# Genome-wide epigenomic stress response in the European sea bass (*Dicentrarchus labrax*, L.)

**DOI:** 10.1101/2020.06.19.160812

**Authors:** M.V. Krick, E. Desmarais, A. Samaras, E. Gueret, A. Dimitroglou, M. Pavlidis, C.S. Tsigenopoulos, B. Guinand

## Abstract

**Background:** While the stress response inspired genome-wide epigenetic studies in vertebrate models, it remains mostly ignored in fish. We modified the epiGBS (epiGenotyping By sequencing) technique to explore changes in genome-wide cytosine methylation to a repeated acute stress challenge in the nucleated red blood cells (RBCs) of the European sea bass (*Dicentrarchus labrax*). This species is widely studied in both the natural and farmed environments, including issues regarding health and welfare.

**Results:** We retrieved 501,108,033 sequencing reads after trimming, with a mean mapping efficiency of 73.0% (unique best hits). Fifty-seven differentially methylated cytosines (DMCs) close to 51 distinct stress-related genes distributed on 17 of 24 linkage groups (LGs) were detected between RBCs of pre- and post-stress individuals. Literature surveys indicated that thirty-eight of these genes were previously reported as differentially expressed in the brain of zebrafish, most of them involved in stress coping differences. DMC-related genes associated to the Brain Derived Neurotrophic Factor, a protein that favors stress adaptation and fear memory, are especially relevant.

**Conclusion:** We provide an improved epiGBS protocol with increased multiplexing and sequencing capacities that offer new opportunities to improve data acquisition and to investigate important biological processes at a genome-wide level, such as the stress response. Minimally invasive RBCs deserve more attention to investigate the epigenetic response to stress without sacrificing fish.

## Background

The study of stress is of large interest in biological sciences. Stress affects cells as well as ecosystems and influences processes that span from seconds to evolutionary time scales [1,2]. Reaching a common definition is quite complex [2–5], but stress often refers to the non-specific, total response of one organism to answer to internal and/or external stimuli acting as stressors. Accordingly, the stress response represents the repertoire of physiological, metabolic and behavioural changes developed during and observed after a threat to counteract the effect of stressors and recover or approach the homeostatic equilibrium. In teleost fish – the largest group of vertebrates - an integrative understanding of the stress response is still expected [1,2]. This concerns natural populations that may develop plastic and adaptive response to stressors [1,6], but also cultured species in order to increase fish health and welfare in stressful farming environments [7–9].

Classically, stress studies have to integrate physiological, immunological, neurobiological and behavioural components that altogether shape the stress response in fish [8, 10–12]. The stress response implies - among others - a primary neuroendocrine and a secondary immune response, with interacting roles of neuropeptides and cytokines in each respective system [13–15]. The Hypothalamus-Pituitary-Interrenal (HPI) axis plays a crucial role in the manifestation and regulation of the stress response through the release of neuropeptides, catecholamines and glucocorticoids. In teleosts, cortisol is the primary circulating glucocorticoid and species-specific differences in the timing, magnitude and peak duration of post-stress plasma cortisol concentrations have been reported [16]. Glucose - another classical biomarker of stress - is also under control of hypothalamus, and its blood level depends on glycogen metabolism as well as gluconeogenesis, themselves triggered by interrenal glands and catecholamines [17, 18]. Beyond the classical HPI axis, several mediators of allostasis, organs and systems (e.g. serotonin and the serotonergic system; osteocalcin and the parasympathetic nervous system) are involved in stress and anxiety [19–21]. A proper regulation of the stress response requires the consideration of a complex regulatory network of non-linear actions to integrate the whole organism response to stressors [22, 23].

In the past few years, the stress response largely inspired genome-wide epigenetic studies in model organisms [24–28]. For animal models, it was established that stress can especially reshape the stress axis/network through epigenetic modification of genes in the hippocampus, hypothalamus and other stress-responsive brain regions [23,29,30]. Human stress studies have shown that blood cells responded to DNA methylation in the brain [31–33, but see 34], and more generally that blood is useful whenever target tissues cannot be easily sampled or biopsied [35]. Because blood cells are mobilized as part of the stress response in fish [36, 37], it is appealing to investigate if an epigenomic signature of stress is present in their nucleated red blood cells (RBCs). The use of RBCs in fish transcriptomics or proteomics has received attention over the last decade [38–40]. However, fish epigenetics has not embarked on this path yet and genome-wide epigenetic studies using RBCs are quite rare in fish [41]. This contrasts with the over-increasing interest to investigate epigenomic variation in teleosts [41–72] Nevertheless, documenting the epigenomics of the stress axis/network also remains scarce [58], while authors asked for greater attention on this topic [73,74].

After salmonids, the European sea bass (*Dicentrarchus labrax*) is certainly the most investigated marine fish species in Europe using molecular tools. It has been extensively studied over the last three decades, for both natural and farmed populations (reviewed in [75]). This includes the sequencing of its genome [76] and an increasing number of epigenetic studies [77–84]. The stress response of European sea bass remains evaluated using blood parameters that have been shown to reflect multiple components of its stress response [16,85], despite authors have proposed alternatives [86,87]. How the European sea bass methylome may capture such components is actually missing.

We adapted the epiGenotyping By sequencing (epiGBS) protocol originally proposed by Van Gurp et al. [88] to assess the genome-wide epigenomic variation in the RBCs of *D. labrax* submitted to periods of acute stress. EpiGBS targets variation in cytosine methylation – the covalent addition of a methyl group to cytosine nucleotides – that has long been accepted as an important epigenetic modification in many organisms [89,90]. This modification integrates a second restriction enzyme and further multiplexing of individuals, saving costs. Our aims were to explore the epigenomic landscape of the stress response in the European sea bass using this protocol, and to investigate if nucleated RBCs from minimally invasive blood samples could provide relevant information on this issue in an economically important fish like sea bass.

## Results

Nineteen out of 20 sea bass families were represented by at least one individual among the 74 samples analyzed in this study. Fish number per family ranged from one (families A, D, N) to nine (family R) individuals. Except for the families with a single representative and family M with post-stress fish only (four), both pre-stress and post-stress individuals were present in the 15 remaining families. Four individuals from four distinct families were retained twice by chance, and were thus analyzed for both pre- and post-challenge stress conditions. A total of 70 distinct fish has been analyzed in this study.

### EpiGBS library construction and sequencing

We obtained 504,271,331 total sequencing reads of which 99.4% (501,108,033) were retrieved after trimming of our single library. After demultiplexing, read numbers per sample ranged from 2,284,915 to 16,314,759, with an average of 5,212,596 reads per sample (see Additional File 1). Demultiplexed samples were mapped against the *D. labrax* reference genome with a mean mapping efficiency of 74.5% (73.0% for unique best hits; Additional File 1). Sequencing reads mapped across all linkage groups (Fig. 1). The mean per base pair read depth was 250X.

**Figure 1:**
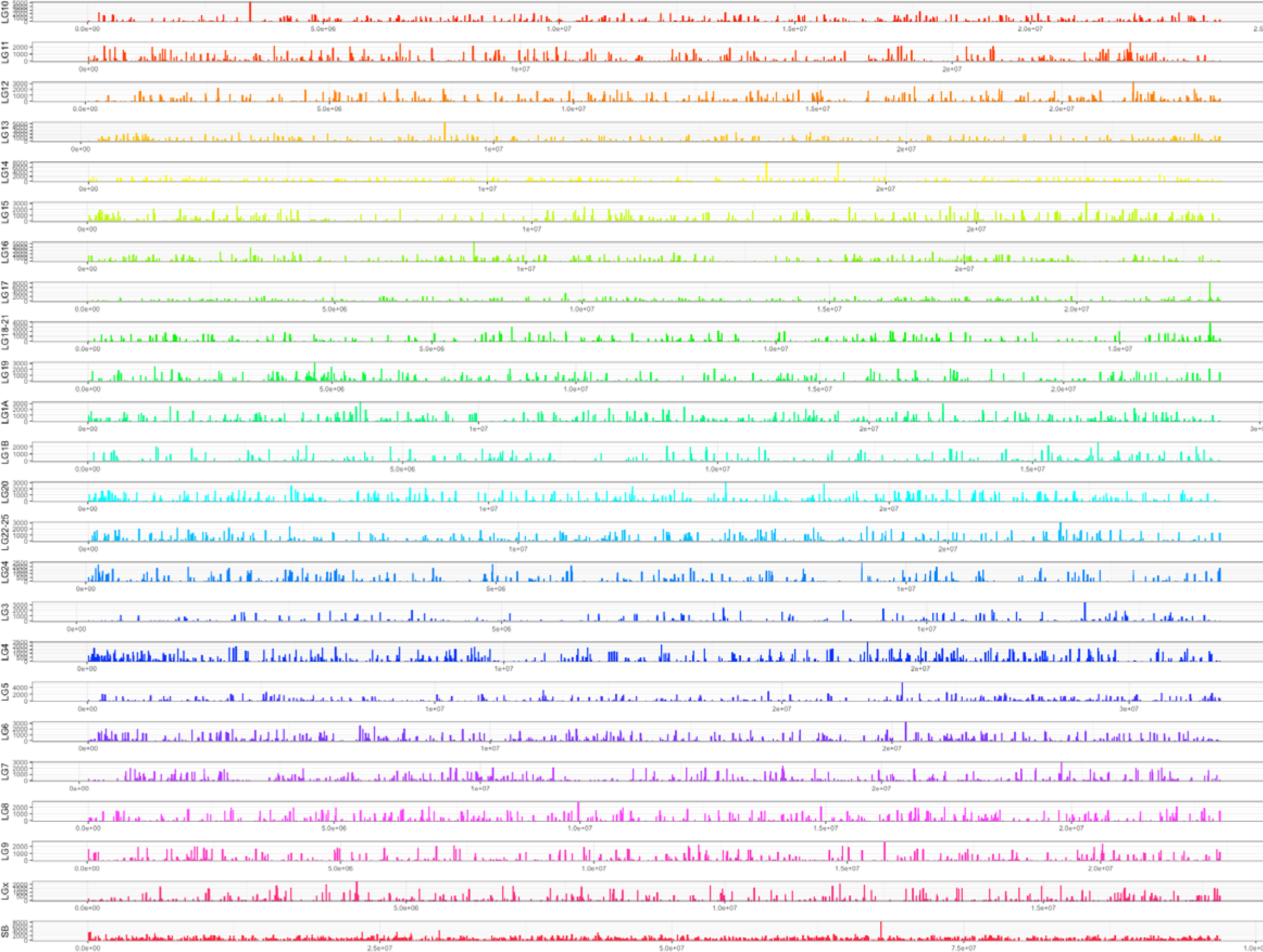
Coverage of the 24 linkage groups (LGs) of the sea bass genome by sequencing epiGBS reads. Ordinates represent the mean number of sequencing reads. LG numbers are those used in [76]. Complex IDs resulting from the merging of previously defined LGs during final assembly (e.g. LG22-25 resulted in the fusion of previous LG22 and LG25). SB-UN (sea bass – unknown; hereby SB) indicates a composite LG made from originally unassembled scaffolds [76].

### Methylation analysis

Out of the 10,368,945 CG dinucleotides (0.46% of the whole) present in the *MspI-SbfI* reduced-representation of *D. labrax* genome we obtained, 47,983 CpG coordinates were extracted with a minimum of 30X read depth and presence in at least 20 individuals. They were filtered out using a 15% methylation difference threshold and a nominal cut-off value of *q* < 0.001. With these parameters, only a total of 57 cytosines in CpG context were defined as DMCs between pre- and post-stress sea bass (Table 1). Methylation differences ranged up to 46.4% for hypermethylated cytosines, and down to −27.5% for hypomethylated cytosines. Hyper-methylation was more frequently detected than hypomethylation (11 [19.30%] hypo- *vs* 46 [80.70%] hypermethylated DMCs) in post-stress sea bass. DMCs were distributed on 17 out of 24 LG groups and in or close to 51 distinct genes. Further information is provided in Additional File 2 (e.g. gene annotations, CpG context).

**Table 1:** Differentially methylated cytosines (DMCs) found in this study between pre- and post-stress sea bass

| *D. labrax* LG | Ensembl LG | q-value | methylation difference (%) | *D. labrax* Name | Gene ID | Gene Name | Location |
| --- | --- | --- | --- | --- | --- | --- | --- |
| **LG1A** | **HG916837.1** | **0** | **-21,40** | **DLAgn_00094580** | **FOXJ3** | **forkhead box protein J3** | **gb** |
| **LG1A** | **HG916837.1** | **0** | **-21,67** | - | - | - | **gb** |
| **LG1A** | **HG916837.1** | **0** | **-15,51** | - | - | - | **gb** |
| LG3 | HG916843.1 | 1,54E-95 | 18,14 | DLAgn_00141440 | ABLM2 | actin binding LIM protein family -- member 2 | gb |
| LG4 | HG916844.1 | 1,08E-84 | 16,41 | DLAgn_00145290 | CELA3b | proproteinase E-like (elastase 2) | gb |
| LG4 | HG916844.1 | 0 | 21,38 | DLAgn_00146200 | NCBP2 | nuclear cap-binding protein subunit 2 | int. |
| LG5 | HG916845.1 | 3,67E-265 | 21,91 | DLAgn_00159160 | CILP1 | cartilage intermediate layer protein 1 | gb |
| LG5 | HG916845.1 | 8,93E-89 | 16,64 | DLAgn_00159900 | GRPT1 | growth hormone regulated TBC protein 1 | gb |
| LG6 | HG916846.1 | 6,44E-245 | 20,44 | DLAgn_00164450 | GLG1a | golgi glycoprotein 1 | gb |
| LG6 | HG916846.1 | 0 | -21,01 | DLAgn_00166360 | GDPGP1 | gdp-d-glucose phosphorylase 1 | exon |
| **LG6** | **HG916846.1** | **0** | **-18,98** | **DLAgn_00172370** | **KBTBD13** | **kelch repeat and btb domain-containing protein 13-like** | **int.** |
| **LG6** | **HG916846.1** | **0** | **-18,18** | - | - | - | **int.** |
| **LG6** | **HG916846.1** | **0** | **-16,12** | - | - | - | **int.** |
| LG8 | HG916848.1 | 2,77E-247 | 22,52 | DLAgn_00186250 | BTPF | nucleosome-remodeling factor subunit bptf | gb |
| LG10 | HG916827.1 | 2,44E-79 | 15,44 | DLAgn_00005890 | PBX1 | pre-B-cell leukemia homeobox 1 | gb |
| LG10 | HG916827.1 | 6,98E-125 | 21,55 | DLAgn_00006210 | ADCY1 | adenylate cyclase 1 (brain) | gb |
| LG11 | HG916828.1 | 0 | 27,70 | DLAgn_00016970 | MMNR2 | multimerin 2a Precursor Elastin Microfibril Interface Located Elastin Microfibril Interfacer | gb |
| LG12 | HG916829.1 | 0 | 22,32 | DLAgn_00019700 | FBXO33 | F-box protein 33 | int. |
| LG12 | HG916829.1 | 1,23E-128 | 20,01 | DLAgn_00025280 | SASH1A | sam and sh3 domain-containing protein 1-like | int. |
| LG14 | HG916831.1 | 1,48E-53 | 15,74 | DLAgn_00037440 | COL4A5 | collagen type IV alpha 5 chain | int. |
| LG14 | HG916831.1 | 0 | 42,05 | DLAgn_00037440 | COL4A5 | collagen type IV alpha 5 chain | gb |
| LG14 | HG916831.1 | 0 | 22,46 | DLAgn_00038760 | ROBO3 | roundabout homolog 2-like | int. |
| LG14 | HG916831.1 | 5,22E-212 | 17,93 | DLAgn_00044110 | CLDN4 | claudin 4 | gb |
| LG14 | HG916831.1 | 1,25E-155 | 27,48 | DLAgn_00046110 | TMEM132E | transmembrane protein 132e | gb |
| LG16 | HG916833.1 | 2,78E-168 | 26,24 | DLAgn_00063590 | CSMD3a | CUB and Sushi multiple domains 3a | gb |
| LG16 | HG916833.1 | 9,78E-92 | 17,12 | DLAgn_00063770 | TRMT11 | tRNA methyltransferase 11 homolog | int. |
| LG16 | HG916833.1 | 8,81E-215 | 17,80 | DLAgn_00064610 | FURIN | furin-like protease kpc-1 | gb |
| LG16 | HG916833.1 | 0 | 17,76 | DLAgn_00064820 | SPIRE1b | protein spire homolog 1-like | gb |
| LG17 | HG916834.1 | 1,91E-82 | 18,84 | DLAgn_00073160 ;<br>DLAgn_00073170 | PLG (sense);<br>SLC22A2 (antisense) | Plasminogen (sense) / Solute carrier family 22 member 2-like (antisense) | gb / 3'UTR |
| LG17 | HG916834.1 | 1,28E-231 | 19,86 | DLAgn_00073770 | RRM2 | ribonucleotide reductase regulatory subunit M2 | gb |
| LG20 | HG916840.1 | 2,76E-207 | 15,01 | DLAgn_00114460 | TNKSb | tankyrase, TRF1-interacting ankyrin-related ADP-ribose polymerase b | gb |
| LG20 | HG916840.1 | 1,34E-115 | 15,35 | DLAgn_00114810 | LRRTM4L2 | leucine-rich repeat transmembrane neuronal protein 4-like | gb |
| LG20 | HG916840.1 | 3,78E-68 | 15,32 | DLAgn_00116160 ;<br>DLAgn_00116150 | FICD (sense) ;<br>SART3 (antisense) | Adenosine monophosphate-protein transferase ficd-like (sense) / Spliceosome associated factor 3, U4/U6 recycling protein (antisense) | gb |
| LG20 | HG916840.1 | 1,02E-88 | 18,04 | DLAgn_00121270 |  | Coding region of a truncated Non LTR Retrotransposable Element (RTE) RET-1_AFC | rr |
| LG20 | HG916840.1 | 0 | 35,61 | DLAgn_00122640 | BMP3 | Bone morphogenetic protein 3 | gb |
| LG20 | HG916840.1 | 0 | 15,40 | DLAgn_00124110 | ZMAT4 | zinc finger matrin-type 4b | gb |
| LG22-25 | HG916841.1 | 7,62E-96 | 16,89 | DLAgn_00125750 | NOL4LB | nucleolar protein 4-like b | gb |
| LG22-25 | HG916841.1 | 3,26E-99 | 15,05 | DLAgn_00132500 | CCDC30 | coiled-coil domain containing protein 30 like | gb |
| LG24 | HG916842.1 | 2,81E-43 | -15,63 | DLAgn_00136190 | GLI2 | zinc finger protein gli2-like | gb |
| LG24 | HG916842.1 | 2,87E-112 | 16,47 | DLAgn_00136980 | LRRC3 | leucine rich repeat containing 3 | int. |
| LG24 | HG916842.1 | 0 | 30,68 | DLAgn_00137190 | UNC80 | Protein unc-80 homolog isoform 2 | gb |
| LG24 | HG916842.1 | 0 | 15,42 | DLAgn_00137560 | CHN1 | N-chimerin | gb |
| LG24 | HG916842.1 | 9,47E-204 | 17,08 | DLAgn_00139980 | PTGFRN | prostaglandin f2 receptor negative regulator | gb |
| LGx | HG916850.1 | 4,18E-251 | 19,94 | DLAgn_00209310 | MYF5 | myogenic factor 5 | gb |
| LGx | HG916850.1 | 4,52E-204 | 23,46 | DLAgn_00209760 | CELF2 | cugbp elav-like family member 2 | gb |
| SB-UN | HG916851.1 | 1,61E-152 | 16,29 | DLAgn_00218300 | PRKCQ | protein kinase c theta type | int. |
| SB-UN | HG916851.1 | 1,09E-120 | 16,34 | DLAgn_00220000 | PTPRB | protein tyrosine phosphatase receptor type B | gb |
| SB-UN | HG916851.1 | 6,07E-154 | 18,95 | DLAgn_00222790 | NPAS3 | neuronal PAS domain-containing protein 3-like | gb |
| **SB-UN** | **HG916851.1** | **0** | **-21,55** | **DLAgn_00227120** | **CRTC2** | **CREB regulated transcription coactivator 2** | **gb** |
| **SB-UN** | **HG916851.1** | **6,11E-56** | **15,79** | - | - | - | **gb** |
| SB-UN | HG916851.1 | 4,15E-61 | 17,53 | DLAgn_00227130 | DENND4B | Denn domain-containing protein 4b | gb |
| SB-UN | HG916851.1 | 1,51E-46 | 15,84 | DLAgn_00236500 | GFRA2 | GDNF family receptor alpha-2 | gb |
| SB-UN | HG916851.1 | 0 | 46,39 | DLAgn_00238280 | DLG1 | disks large homolog 1-like | gb |
| **SB-UN** | **HG916851.1** | **9,43E-157** | **26,15** | **DLAgn_00242570** | **BTR30** | **E3 ubiquitin-protein ligase TRIM39-like (bloodthirsty-related gene family, member 30)** | **gb** |
| **SB-UN** | **HG916851.1** | **6,99E-209** | **-17,71** | - | - | - | **gb** |
| SB-UN | HG916851.1 | 2,11E-216 | -27,49 | DLAgn_00244380 | GMPPB | GDP-mannose pyrophosphorylase B | exon |
| SB-UN | HG916851.1 | 5,09E-256 | 20,97 | DLAgn_00245980 | MATR3 | matrin-3 | exon |

Location on the European sea bass linkage groups (LGs) of the 57 differentially methylated cytosines (DMCs) found in this study between pre- and post-stress individuals. The four differentially methylated regions (DMRs) are in bold. For each DMC, the false-discovery rate adjusted q-values at the nominal $q = 0.001$ cut-off threshold are reported, together with their methylation difference. Gene names and gene symbols (IDs) of DMC-related genes ( $n = 51$ ) are reported. The location of each DMC is given (gb: gene body, int.: intergenic, 3'UTR or rr: repeat region). We did not arbitrarily defined promoter regions in this study. An extended version of this table reporting annotations and further useful information are offered in Additional File 2.

Most identified DMCs were found located within identified gene bodies (44 out of 57, 77.19%), one in the 3’UTR regions of the Solute Carrier family 22 Member 2 (*SLC22A2*) gene on LG17, and one in a repeated region (a non LTR Retrotransposon Element on LG20). In the remaining cases (*n* = 11), DMCs are intergenic and located in a window ranging from 0.9kb to 51kb to the closest gene (respectively: *SASH1A* on LG12 and *TRMT11* on LG16; Table 1). Two pairs of overlapping, but inversely oriented genes share on their sense vs. antisense strand an identical DMC: *PLG* and *SLC22A2* on LG17, and *SART3* and *FICD* on LG20 (Table 1)

When located on the same LG, DMCs were usually distant by at least 30kb from each other. In only four instances, DMCs occurred as differentially methylated regions (DMRs) that grouped 2-3 DMCs separated by only few hundreds of base pairs (Table 1). This includes three hypo-methylated cytosines grouped in a DMR located ~1500 bp downstream of the predicted Kelch Repeat and BDB domain 13 (*KBTBD13*) gene with at most 88 bp between the cytosines. Another DMR includes three hypomethylated cytosines (>20%) on LG1A in the second intron of the forkhead box J3 (*FOXJ3*) gene. Two other groups of two cytosines were found either in the same exon (distant by 3 bp) of the *BTR30* gene or in two different introns (distant by 3.8Kb) of a predicted CREB regulated transcription coactivator 2 (*CRTC2* or *TORC2:* Transducer Of Regulated cAMP Response Element-Binding Protein [CREB] 2) (Table 1). In these two latter cases, one cytosine was found hypomethylated while the other is hypermethylated. A single DMC was associated to a repeat region and two DMCs were found to refer to the same gene (homologous to the *Gasterosteus aculeatus* paralogue of *COL4A5*, a collagen gene of type IV mostly implicated in the protein network of the basement membrane) (Table 1). For this gene, one DMC is located in the first intron while the second is 16.5kb upstream of the start codon. They were not considered as a DMR.

### Clustering

Hierarchical clustering showed a strong family effect in methylation patterns (i.e. individuals within family clustered together; Fig. 3). The four individuals that were caught twice clustered together by pairs in all four cases. These individuals have the lowest levels of dissimilarity in hierarchical clustering, suggesting very few differences in their methylation profiles in pre- *vs* post-stress condition. Despite this strong family effect and clues of low impact of the stress on methylation, pre- (T0) and post-stress (T4) groups can be distinguished based on their DMC profile in PCA. Mean loading scores of individuals were found significant among T0 and T4 for PC1 that explained 7.2% of total variance (Student *t*-test; *p* < 0.005, Fig. 4). No significant difference was found for loading scores along PC2 (3.0% of total variation; *P*= 0.404).

**Figure 2:**
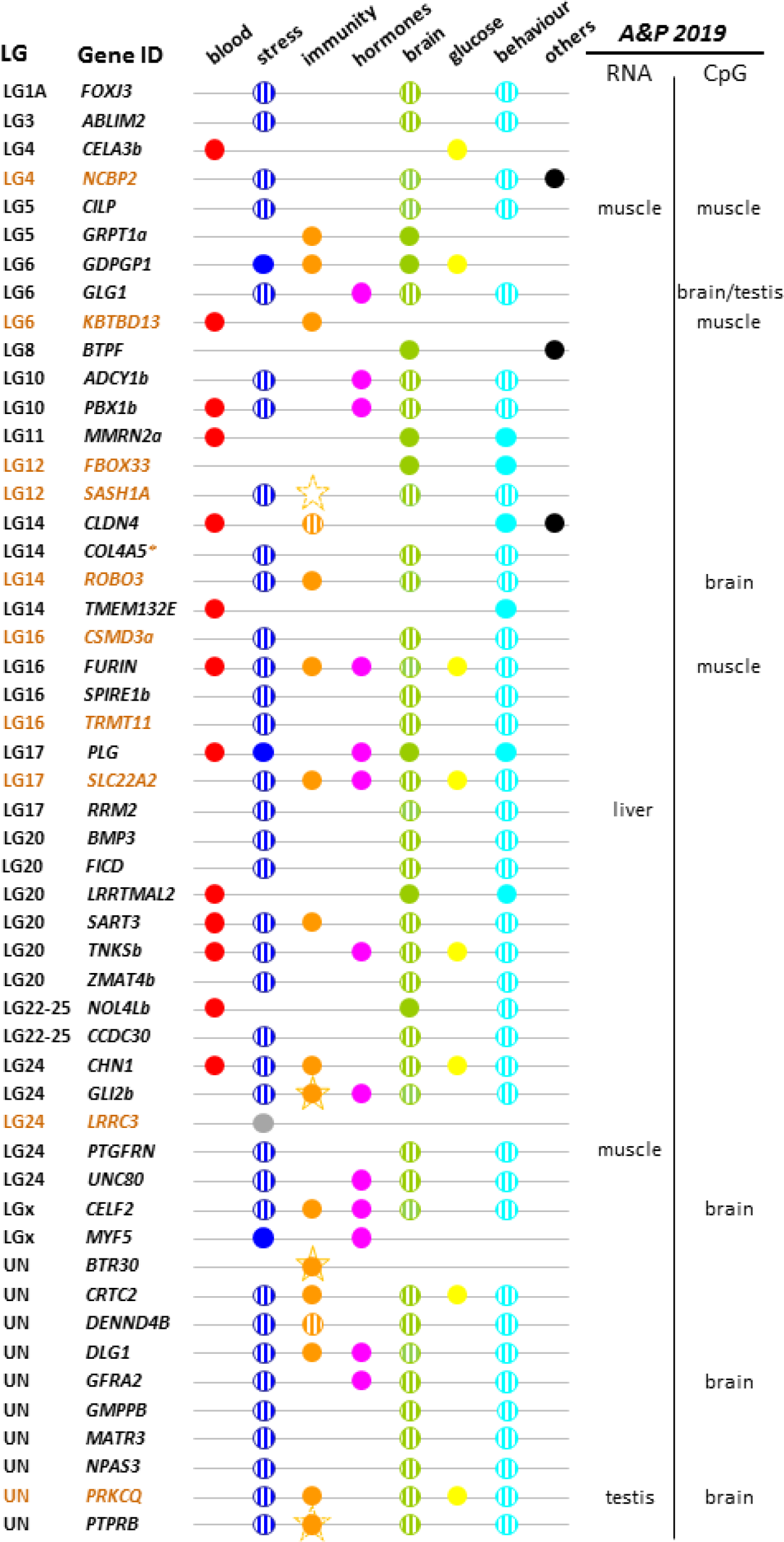
Diagrammatic representation of the involvement of the 51 DMC-related genes found in this study and issues regarding the stress response. One additional issue was added for few genes that are involved in the regulation, the control and the dynamics of epigenetic process (*NCBP2, GTPF*), and acid-base homeostasis in kidney without formal proof of actual relation with a stress issue (*CLDN4*). DMCs located within gene bodies are indicated in black, in brown for intergenic regions including promoters. The hatched symbols refer to genes also reported in high-throughput transcriptomic studies in zebrafish [93–96] (main text and Additional File 3). This does not mean that our literature survey was supported only by these studies, but it illustrates their importance over results. Stars with full line indicate genes detected as differentially expressed (*BTR30, GLI2*) in one immunity-related transcriptomic study in sea bass [91], while stars with broken lines indicate genes with methylation differences (*PTPRB, SASH1A*) (stickleback: [54]; rainbow trout: [58]; respectively). Reports outside of the above-mentioned studies are indicated by filled symbols. On the right are indicated genes found differentially expressed (RNA) or differentially methylated (CpG) in the genome-wide study by Anastasiadi and Piferrer (*A&P 2019*) on European sea bass [83]. Tissues in which genes were found differentially expressed/methylated are indicated (brain not investigated in transcriptomics). Association with the *LRRC3* gene is reported in grey as leucine-rich repeat (LRR) gene family is classically involved in neuroimmunity. Variations in transcription or methylation have been shown to impact behaviours after stress events within this family (see e.g. *LRRTM4L2* in this study), but this relationship remains presumptive for *LRRC3*. Additional support including references for these DMC-related genes is reported in Additional File 3, as well as comparisons with other epigenomic studies.

**Figure 3:**
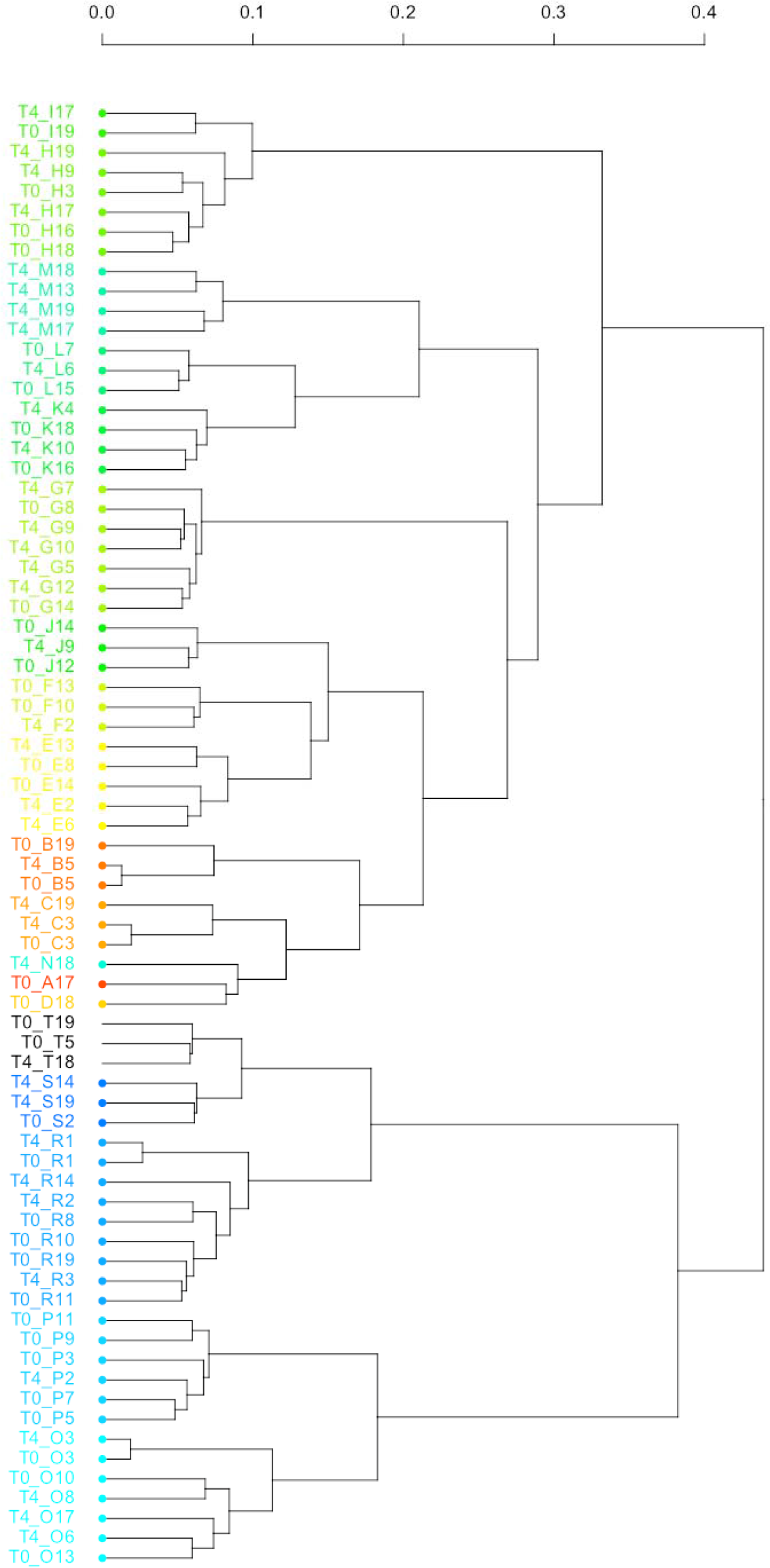
Hierarchical clustering based on of the 57 differentially methylated cytosines (15% threshold) of the randomly caught 74 pre- and post-stress sea bass samples considered during this study. Nineteen out the twenty families (A to T) used in this study are present in this graph and each family is associated to one single colour. Capital letters refer to the family, while T0 and T4 are for pre- and post-stress individuals, respectively.

**Figure 4:**
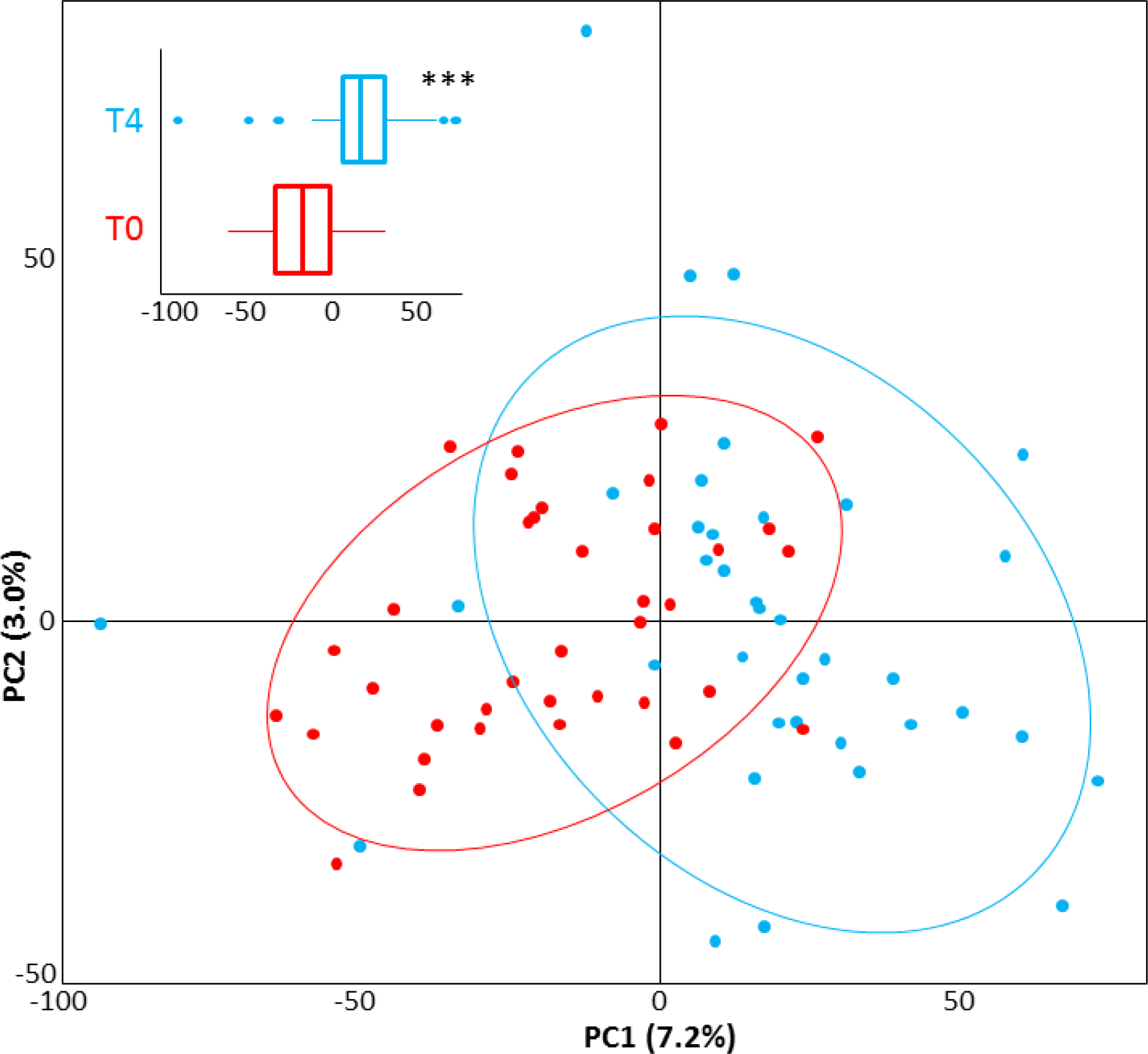
PCA based on the methylation profiles of the 57 differentially methylated cytosines (15% threshold) reported in this study (Table 1). Pre- and post-stress individual sea bass (in red and blue, respectively) differ significantly along PC1 (*p* < 0.001), but not PC2. The insert illustrate the distribution of individual scores along PC1. Ellipses represent the 95% confidence limits over PC1 and PC2.

### Literature surveys

Several literature surveys (see the Methods section for details) were performed to better understand if DMC-related genes detected in sea bass RBCs could effectively (*i*) represent real candidates to investigate the stress response if fish, and (*ii*) be indicative of processes occurring in the brain.

#### Thematic survey

Our first literature survey showed that DMC-related genes detected in this study were found involved in one of the eight themes we retained to describe the stress response in fish (Fig. 2 and Additional File 3). Most of them were previously analysed or detected in human or mice stress studies, in few cases in other model organisms (see Additional File 3). Additionally, the *GLI2* gene encoding for a zinc-finger protein and the *BTR30* (bloodthirsty-related gene family member 30, also known as E3 ubiquitin-protein ligase *TRIM39*-like) gene (Table 1) were found differentially regulated in sea bass cultured-cells submitted to viral infection and thus involved in the immune response [91].

#### Transcriptomics survey in zebrafish

In a second time, we searched for differential expression of sea bass DMC-related genes in studies using high-throughput transcriptomics in fish blood. Only four studies were retained [38–40,92], and none of our DMC-related genes was found to be differentially expressed in these studies.

After curation of PubMed lists and addition of few undetected studies, we surveyed 22 zebrafish studies analyzing the transcriptomics of the stress response in six tissues: brain (*n* = 10), liver (*n* = 7), muscle (*n* = 2), gonads (*n* = 1), intestine (*n* = 1), skin (*n* = 1) (no high-throughput transcriptomics study in gill, kidney, one in muscles of eye that was accounted for ‘muscle’; Additional File 4). The main outcome of this survey was to document 38 out of 51 RBC’s DMC-related genes (74.51%) in stress-related studies. Strikingly, these 38 DMCs were reported mainly from few neurotranscriptomics zebrafish studies that dealt either with reactive-proactive behavioural response to stress [93, 94] or with changes in social regulation that may promote stressful behaviour among congeners [95, 96] (Fig. 2 and Additional File 3). Some genes are shared among these studies. A recent studies recorded five of them as also differentially expressed in Siamese fighting fish (*Betta splendens*) brain during male contests [97] (see Additional File 3), *FBOX33* was not formerly reported in zebrafish, suggesting that 39 DMCs found in sea bass RBCs could be associated to stress.

Only four of these DMC-related genes were reported in the former RNA sequencing study of Anastasiadi and Piferrer [83] in sea bass. This included *CILP1* and *PTGFRN* in muscle, *PRKCQ* in testis, and *RRM2* in liver (blood and brain not investigated) (Fig.2). In comparison, the twelve other zebrafish studies investigating other organs than brain retrieved only 15 of the sea bass DMC-related genes detected in RBCs (see Additional File 4).

#### Epigenomic survey

We finally looked at former studies dealing with epigenomics in fish. The studies retained (*n* = 31) are presented in Additional File 5. Twenty studies reported differential methylation at genes also found differentially methylated in sea bass RBCs (see Additional File 5). Data sets were highly heterogeneous for different reasons (e.g. experimental set-ups, nature of stressors if any, thresholds considered to define a DMC/DMR). Main observations might be summarized as follows: all studies considering brain tissue for which gene annotations were made available (*n* = 6) reported at least one match with DMC-related genes found in the present study. More specifically, data of the single former sea bass epigenetic study performed at a genome-wide level [83] reported five genes of the present study as differentially methylated in the brain (*GLG1a, GFRA2, CELF2, PRKCQ, ROBO3*), three in muscle (*CILP1, FURIN, KBTBD13*), one in testis (*GLG1a*), and any in liver (total: eight distinct genes). This indicated slightly more matches with the brain than with other organs investigated so far. However, the single previous study analyzing RBC methylome in fish so far [41] shared no DMC-related gene with the present study. Finally, the study that reported the largest number of genes also found as DMC-related genes in sea bass (*n* =15) was by Gavery et al. [45]; These authors were interested in the liver of hatchery vs wild rainbow trout during a so-called ‘temporal analysis’ (see Additional File 5). Another study on zebrafish reported 11 genes in common with the present study, but was not organ specific (see Additional File 5) [53].

### Protein-protein interactions

Further database mining screening for specific protein interactions on the String server revealed few other possible pairs of associations (*n* = 8): *ROBO3-CHN1, ROBO3-PRKCQ, ROBO3-LRRC3, DLG1-NCBP2, FURIN-PLG, PLG-MMRN2a, CELF-RRM2*, and *CRTC2-DENND4B* (see Additional File 6). The *FURIN-PLG* and *DLG1-NCBP2* associations were also detected in our literature survey. The first one will be presented in the ‘Discussion’ (see Additional File 3 for the second association). However, most of these associations retrieved only poor information relative to the stress response. *ROBO3* and *CHN1* have been shown to interact in the brain with poorly-understood implications of *CHN1* in neurological disorders [98] (*CHN1* may have other stress-related functions, see Additional File 3). *CELF2* (CUGBP Elav-like family member 2) and *RRM2* (Ribonucleotide reductase M2 polypeptide) are both known to participate to messenger RMA (mRNA) metabolism [99]. *CELF2* acts to post-transcriptionally stabilize mRNAs by relocating them to stress granules in the cytosol. *CELF2* interferes with *RRM2* that modulates its splicing activity. As post-transcriptional activities are at the core of methylation studies, the detection of this association is relevant to our study. To illustrate some limits of these associations: *CRTC2* and *DENN4B* were detected apparently because they are located close to the sex determining region of *Onchorhyncus mykiss* [100]. No relationship with stress is suspected. The relationship between multimerin (*MMRN2*) and plasminogen (*PLG*) via its co-activator remains poorly investigated and supported [101]. This automatic search using String did not really enrich our findings.

## Discussion

We showed that a modified epiGBS protocol originally proposed by Van Gurp et al. [88] was applicable to further analyze patterns of cytosine methylation at a genome-wide scale in *D. labrax*. This is the first use of epiGBS in fish and the second in an animal species (Canadian lynx [102]). The addition of a second restriction enzyme illustrates the flexibility of the epiGBS and more generally of reduced-representation bisulfite sequencing (RRBS) protocols to improve data acquisition and impact [103]. This is however not the first RRBS protocol dedicated to the addition of a second restriction enzyme [104], but this is hereby proposed in a context of improved multiplexing of samples and increased sequencing capacities, thus offering more opportunities to explore important biological processes, such as the stress response. Data allowed us to explore how RBC’s DNA methylation may respond to stressors in the European sea bass and - albeit some limitations discussed below - was shown to complement both the traditional evaluation of the stress response based on analyses of blood serum samples [85,105], or on gene expression variation, often using invasive brain samples [93,94,106–111].

### Mining the epigenome…

The information provided in this study is based on the analysis of 47,983 distinct methylated sites distributed over all sea bass LGs. The mapping efficiency was high (74.5%) when compared to early values retrieved in human (~65%) [112], or in some recent fish studies screening for genome-wide methylation (e.g. 55-60% in [59]; 40% in [41]). Other studies reported similar mapping efficiencies, but reported percentages of mapping for unique best hits that were generally lower. For example, in *Kryptolebias marmoratus*, Berbel-Filho et al. [43] reported a mean mapping efficiency of 74.2% but 61.1% unique best hits while, in this study, this latter percentage reached 73.0%. This reflects a more robust mapping of the DMCs we detected and significantly enlarge the breadth of the sites that can confidently exploit to retrieve functional information. Taking advantage of the epiGBS protocol that allow to process more samples [88], the number of individuals considered in this study is rather high (*n* = 70 distinct individuals), when most epigenomic studies in fish dealt with less than 30 individuals (range: *n* = 3 in [60]; *n* = 106 in [47] for a population study). In sea bass, Anastasiadi and Piferrer [83] previously reported a study that used 27 samples and as many libraries to be sequenced while our data were obtained from a unique library preparation. Our modified epiGBS protocol provides a considerable amount of information, certainly at a reasonable cost, to decipher methylation landscapes of sea bass or other species.

The operational and statistical thresholds used in the successive steps of this study are large, resulting in the discovery of a rather low total number of methylated sites. For example a threshold of 30X and nominal cut-off value of 0.001 are quite conservative, when some studies might consider a threshold of 5X or 10X for a CpG to be analysed and associated cut-off values of 0.05 or 0.01 (e.g. [46,59,83]). Relaxing thresholds would enable to retrieve more DMCs, but elevated thresholds should normally ensure that access to relevant information is reached. Thus, only 57 DMCs have been found in RBCs of pre- and post-stress European sea bass, and only few were grouped as DMRs. Considering DMRs rather than DMCs is common in genome-wide methylation studies, including sea bass [83]. Methylation changes grouped in clusters (i.e. DMRs) are more likely to influence transcriptional activity at nearby loci than single cytosines [113]. However, other experimental issues associated to the flexibility of RRBS protocols (e.g. single- *vs* paired-end sequencing, likelihood of annotation, selection and size of fragments) can made the advantages of considering DMRs over DMCs perhaps more disputable [103] and DMRs are not always accessible [114]. In this study, DMCs were found mostly hypermethylated in post-stress individuals compared to the pre-stress individuals, and mostly located in gene bodies of fifty-one different genes (44 out of 57; 77.2%). It has been shown that such location of differential methylation may regulate splicing and/or act as alternative promoters to reshape gene expression [115–117]. Thus, despite a low number of DMRs in this study, a large fraction of DMCs points to the transcriptionally active portion of the genome and to important stress-related genes.

In addition to DMCs located in gene bodies, a dozen of DMCs were found in intergenic regions (21.0%). Intergenic cytosine methylation has been frequently described, including in stress studies [118], but its role remains poorly understood [119,120]. While numbers of genic vs intergenic DMCs may greatly vary, a ratio of ~80% of DMCs located in gene bodies and ~20% located in other genomic regions has been reported in other fish studies (e.g. [46]). Sometimes proportions are approximately reversed (~80% intergenic vs ~20% in gene bodies; e.g. [50]). While under-represented in our data, the role for differentially methylated transposable elements is also poorly understood [121, 122], while crucial in the stress response [123].

### … but mining the unknown

Studies looking at the epigenomic landscape of RBCs in fish are scarce. Gavery et al. [41] reported 85 DMRs between hatchery and wild fish in *O. mykiss*. This study suggested that blood DNA methylation patterns are indicative of the stressors experimented in the environment, but we are still currently mining the unknown to gain supporting information.

We nevertheless demonstrate that fish RBCs are relevant candidates to investigate the stress response at the epigenomic level in the European sea bass. Numerous DMC-related genes reported here have been previously shown to respond to stress or implicated in variation of the stress response in animal models. Overall, our comparative surveys showed that the DMC-related genes we detected in sea bass using RBCs have more connections with results found in the brain than in other tissues. This relationship is however weak at the epigenomic level and about one third of our DMC-related genes have also been detected in liver of trout [45]. This relationship is far more pronounced when genome-wide gene expression difference is considered. Indeed, genome-wide neurotranscriptomics studies dealing with personality and stress coping styles [93,94], or social regulation [95] collectively reported thirty-eight of our 51 DMC-related genes as expressed and potentially associated to the stress response in zebrafish (one added when including a study on *B. splendens* [96]). As differential methylation is classically interpreted as leading to differential gene expression (but see, e.g., [124, 125] for a more nuanced consideration), this observation is one interesting signal indicating that RBCs might monitor events occurring in the brain. However, it remains unclear why most of them are related to coping and personality, and to studies where fish were under stress for few minutes [93,94], hours or days [95]. How distinct stress contexts and challenges may apparently rely on similar stress circuitry has to be investigated further.

### A brain-derived neutrophic factor (BDNF) network

The roles of the DMC-related genes have to be exposed more extensively to illustrate their possible involvement in the stress response of sea bass. This is not place to fully expand on all DMC-related genes (see Additional File 3 for details), but one single selected example can be developed. The best representative is a set of DMC-related genes from the BDNF network whose role in the stress response is commonly reported. BDNF is a protein synthesized in the brain that offers resistance to neurodegenerative diseases and favors stress adaptation and resilience [126,127], but also energy homeostasis [128]. BDNF has emerged as one of the most important molecule in memory [129, 130]. It consolidates both the within- and between-generation fear memory owing to epigenetic regulation [131,132]. Its activity is strongly linked to glucocorticoid stress to imprint neurogenesis [133] and it acts as both a regulator and a target of stress hormone signaling [127]. In fish, cortisol binds the glucocorticoid receptor (*gr*) and controls BDNF expression in the brain. With *gr* and relatively few other genes, BDNF has thus emerged as one of the target genes of interest in stress studies, as shown for zebrafish [134], sea bream [135] and sea bass [86,136–138]. BDNF is also involved in other aspects of the stress response in fish (e.g. regulation of the microbiota-gut-brain axis [139]). Variation in BDNF methylation status could not be investigated in this study as no *SbfI* restriction site is present within or close to this gene. Nevertheless, a methylation signature of stress associated to BDNF is likely present in our data and illustrates how blood methylation may provide a snapshot of events studied in the brain.

Indeed, several DMC-related gees are involved in the maturation of proBDNF to mature BDNF or the regulation of its activities (*ABLIM2, ADCY1b, CRTC2, FURIN, NPAS3, PLG*, and possibly *SLC22A2*). Most of these DMC-related genes were already found differentially expressed in the zebrafish brain [93–95], but an outline of their interaction was not presented. The adenyl(ate) cyclase (AC, *ADCY1b* gene) is a brain-specific signaling enzyme that synthesizes the cyclic AMP [140]. In the brain, AC is activated by PACAP (Protein Adenylate Cyclase Activating Protein) and stimulates memory and long term potentiation [141]. This inducible signaling pathway participates to the synthesis of the active form of BDNF (proBDNF to mature BDNF) [142]. ProBDNF is processed by furin and the plasminogen system [128], including processing steps that necessitate actions of actin-binding LIM kinases (ABLIM) [143]. Furin - encoded by *FURIN* - is a subtilisin-like protein proconvertase ubiquitously expressed in fish [144]. It presents an intracellular cleaving activity of numerous biologically relevant molecules in the trans-Golgi network. This includes BDNF in the brain, notably in pre-synaptic axons. In mice, furin has been shown to participate to dendrite morphogenesis and modulate learning abilities and memory [145]. Stress imprinting at *FURIN* is likely and it has recently been shown that transgenerational epigenetic effects of furin activity were active in brain of mice [146]. ProBDNF cleavage by furin depends on brain AC and CREB (cAMP response element-binding protein) signaling [147,148] and plasminogen (*PLG*) [149]. This activity is modulated by stress hormones (corticosteroids) and is essential to brain hippocampal plasticity in mammals [22,150]. One interesting supplementary observation is that CREB signaling necessary to furin is associated to *CRTC2* - a CREB co-activator – found as one of the four DMRs reported in this study. In mice, *CRTC2* is known to act as a switch for BDNF and glucocorticoids to direct the expression of corticotropin-releasing hormone (CRH) in the hypothalamus [151].

Addiitonally, in the brain, plasminogen encoded by *PLG* is converted to plasmin that cleaves BDNF in the extracellular synaptic domain [128,152]. *PLG* has also been shown to regulate pro-opiomelanocortin (POMC) in the hypothalamic-pituitary axis, then the production of peptides hormones such as the adrenocorticotropic hormone (ACTH) [153].

In relation to BDNF, two other DMC-related genes should be mentioned: *NPAS3* (neuronal PAS domain containing protein 3) and *SLC22A2. SLC22A2* - also known as *OCT2* (organic cation transporter 2) – associated to the unique 3’UTR DMC found in this study is implied in numerous transmembrane transports [154], including at the blood-brain barrier [155]. It was found involved in memory in mice [156,157] or *Drosophila* [158], and associated itself to neurodegenerative diseases [159]. Substrates of *SLC22A2* include neurotransmitters such as norepinephrine, but also dopamine, serotonin and a wide-variety of internal or exogenous compounds [160,161]. Functional links of *SLC22A2* with BDNF remain however poorly documented [162], while they have been found co-expressed and co-regulated in the brain during drug administration experiments (e.g. methamphetamine, mimicking the action of catecholamines) [163]. In this study, *PLG* and *SLC22A2* are associated to the same DMC; the functional significance of this situation should be investigated further. Finally, *NPAS3* has a well-established action in memory [164,165]. It participates to neurogenesis in the hippocampus within a network that also includes BDNF [166]. *NPAS3* is also associated to the glial cell line-derived neurotrophic factor (GDNF) receptor-alpha2 gene (*GFRA2*) detected in this study and related to stress and anxiety [167]. The *DLG1* (Disk-large homolog 1) gene could be also associated to BDNF, but this will be not develop further (see Additional File 3).

Overall, results suggest that even in absence of BDNF sequencing reads, some DMCs/DMRs with a potential impact in fear memory and anxiety response as expected in fish after an acute stress challenge certainly did not occur only by chance in this study. They are also related to processes that regulate the HPI axis and hormones. The measurement of methylation difference now used to monitor the BDNF activity and its relationship to stress [132,168] could thus potentially depend on the methylation status observed at or close to *ADCY1b, ABLIM2, FURIN, CRTC2, PLG* and other genes such as *SLC22A2* and *NPAS3*. This should be investigated further to improve our understanding of changes in the directionality of multigenic epigenetic modifications and inter-individual variation in stress coping. Candidate genes coming from other fish or more generally vertebrate studies with role in memory consolidation could be added (e.g. [112,169] for proposals).

As at least *ADCY1b, FURIN, CRTC2, PLG* and *SLC22A2* have other well-recognized relationships with stress and the stress response (especially the glucose metabolism and the immune function; see Additional File 3), this also illustrates the complex non-linear network modulating the stress response in fish with a pivotal role in the brain associated to BDNF and secondary additional outcomes in other tissues and organs.

### Family effect, development and few individuals

While the PCA showed that pre- and post-stress individuals have in average distinct methylation profiles that may represent stress imprinting during the challenge test, a strong family effect was also detected. Family effects were already reported in fish epigenetic studies [51,54,63], and sea bass families certainly do not have the same sensitivity to stress. This may suggest that (*i*) their genomic backgrounds alter how family will develop an epigenetic stress response over the period covered by the challenge test, (*ii*) that the developmental period during which families were distributed over different tanks was a “critical time window” [170] sufficient to imprint individuals and lead them to develop a familial response to stress during the challenge, or that (*iii*) some degree of transgenerational inheritance in methylation profiles is present. These proposals are not mutually exclusive. Transgenerational inheritance has gained interest over the last years [171], notably in marine species [50,172,173], and including sea bass [83]. Parents and/or the germline were unfortunately not sampled in this study for further analysis. As Gavery et al. [41] reported that *BTR30, DLG1, MMRN2*, and *NOL4B* - also found in this study - were differentially methylated in the sperm of *O. mykiss*, these differentially methylated genes could be interesting candidates for transgenerational inheritance study of the stress response. Other candidates are possible. For example, Ghalambor et al. [108] reported rapid evolution in brain gene expression of the Trinidadian guppy (*Poecilia reticulata*) for the *TNKS* and *NCBP2* genes, found as DMC-related genes in this study. It would thus be interesting to investigate how gene expression and cytosine methylation at these genes may co-vary within and between species over generations.

Anastasiadi and Piferrer [83] showed that the domestication process in *D. labrax* resulted notably in genome-wide methylation differences for genes involved in the nervous system and neural crest cell differentiation. In our study, *GLI2* [174], *PBX1* [175], *DLG1* [176] and *FOXJ3* [177] are DMC-related genes involved in neural crest cell proliferation and differentiation. *ROBO3* is involved in the immune response, but also implied in early neurogenesis [178], as well as *NPAS3* [179]. With *BMP3* [180], *FURIN* [181]; *NOL4B* [182], and *Myf5* [183,184], most of these genes are engaged in the development of the anterior region and/or the craniofacial skeleton which is modified during sea bass farming [83]. This suggests that methylation patterns observed in sea bass could also potentially rely on the ontogenetic regulation of a particular phenotype between pre- and post-stress fish, rather than being directly related to the stress challenge. Variation in gene expression or methylation accompanying ontogeny are reported in fish [45, 185] and, as for other organisms, cytosine methylation is known to vary with age in sea bass [84]. Collateral effects of stress might thus be present in our data as suggested by the fact that fifteen genes we found differentially methylated in RBCs were reported as also differentially methylated in the liver of the rainbow trout [45] (see Additional File 5).

On another hand, the epigenetic profiles of four individuals were analyzed in the pre- and post-stress situations and clustered very closely from each other in each case. While based on few observations, this suggests that neither the stress challenge itself, nor development have a strong impact on the cytosine methylation landscape in sea bass. How a transgenerational and within-generation stress-imprinting events influenced by ontogeny may interact to shape both the plastic and the heritable component of the stress response in relation to environmental stimuli require in depth evaluation [186,187]. Nature and strength of family-based epigenomic variation are of considerable importance for selection breeding improvements in a cultured fish like sea bass [188], including also issues about health and welfare [189].

### Blood and RBCs: extending the perspective

As fish engaged in aquaculture programs as sea bass are costly and based on breeders that cannot be easily sacrificed, epiGBS using blood samples may offer a genome-wide assessment of stress-induced epigenetic marks over a significant number of individuals and at reasonable cost. Blood is widely used in human epigenomics, but not in fish despite the presence of nucleated cells and minimally invasive sampling that allows for multiple measurements. Using RNA-sequencing, authors [38,39] already suggested that fish erythrocytes would be useful to provide insights on the innate immunity or response to pathogens. Our study suggests that blood could effectively offer an in depth insight for the evaluation of the stress response from a novel perspective, accompanied by classical physiological measurements (e.g. cortisol, glucose, lactate) and may provide a more complete picture of the stress response in fish than currently performed.

The link between brain and blood epigenomics remains however to be explored more deeply in fish and requires careful evaluation and validation to correct for tissue specificity, as requested in human [190]. Our attempts to survey fish studies undoubtedly showed that considerably more knowledge has to be accumulated before to reach this issue. In human, the lack of access to brain tissues have been shown to impede some epigenomic studies and naturally raised ethical issues certainly also valid for fish [191,192]. *In situ* hybridization techniques in brain have been shown to be especially important to link neurobiological activities and stress coping in fish [134,193]. Their use and thus sacrifice of individuals cannot be ruled out. Actually, methylation differences among full brain or brain areas have been rarely studied in fish, mostly in zebrafish [194], but recently in a tilapia [48], a goby [56], and a killifish [43].

## Conclusion

The European sea bass has become one of the most studied species in fish epigenetics [77–84]. Because of the availability of its genome and the development of selection programs, efforts should be dedicated to further explore its epigenomic landscape regarding the stress response, especially in an aquaculture context. This should be done over a multigenerational scale to identify pathways structuring the stress response at key life stages, on different tissues to better delineate stress-responsive cellular networks, and investigating the effects of the nature, the duration, and the number of stressors. Studies in epigenomics including phenotypic information [195,196] could increase the panel of diagnostic tools for breeding selection in aquaculture if blood could be – at least partly - substituted to brain or other tissues when studying the stress response. Screening for genome-wide epigenomic variation in natural populations as already done for genomic variation [197] is now also conceivable in this species. Our modified version of the original epiGBS protocol seems to be a powerful and affordable method to screen a significant number of individuals with sufficient depth and coverage to reach meaningful conclusions. While it should be compared to others (e.g. [104,198,199]), our protocol is close to the double digestion restriction site DNA sequencing protocol used in genomics. Their parallel use certainly deserves attention to design more integrated epigenomic-genomic studies [200,201], and, more generally, multi-omics investigations of stress, health and welfare [202,203].

## Methods

### Rearing, stress challenge, and blood sampling

Four hundred European sea bass were initially used in this study. Fish were produced in the hatchery facility of Nireus S.A. from breeders maintained in this company for scientific purposes. They resulted of a single crossing experiment (12 dams, 20 sires) that took place in January 2018. During the larval phase, 20 different families labelled from A to T were raised. Each family was reared separately in open circulation tanks at Nireus S.A. research facilities (Greece). To avoid environmental effects, fish were tagged at ~280 days post-hatch (July 10-13^th^ 2018) and distributed in 20 tanks; each tank receiving 1 fish from each family (i.e., 20 fish per tank). Fish were fed twice a day, for 6 days a week, using a commercial diet (Blue Line 45:20 3.5 mm, Feedus S.A., Greece). Throughout the experimental period, the photoperiod was set at 12L:12D, the water temperature and the salinity held constant (18.1 ± 0.2°C and 28 ppt, respectively). Fish weights (mean ± SD) were 48.1 ± 12.8g and 86.2 ± 24.1g in pre- and post-stress individuals, respectively.

Fish were submitted to one acute challenge test per month for three consecutive months, from July to October 2018. During this challenge, fish were exposed to high density stress by lowering water levels in the tank to 1/3 of the original volume, followed by chasing of the fish with a net for five minutes and a 30 min waiting period before sampling. These stressors are classical in sea bass studies regarding response to acute stress [16]. This protocol took place in each tank then for each fish individual entering the experiment. It was repeated for 3 consecutive times at 20-21 day intervals (period long enough for fish to recover).

An initial blood sampling occurred two weeks prior to the implementation of the stress challenge (hereafter T0, July 19^th^, 2018, pre-stress/control group) then at the end of the challenge test (hereafter T4, October 5^th^, 2018). Blood samplings were performed in anesthetized fish. Specifically, fish were anesthetized in 2-phenoxyethanol (300 ppm; Merck; 807291; USA). Recovery was performed in a separate tank with provision of air before fish returned to their holding tank. Blood from the caudal vessel was collected. Plasma and RBCs were separated by centrifugation at 2,000g for 10 min. Careful separation of the plasma and RBCs was performed using 200 μl pipettes. RBC extracts were heparinized (heparin sodium; Sigma-Aldrich), transferred in microtubes and conserved at −20°C in 1 ml of RNA later. DNA was extracted using the Macherey Nagel Nucleo Spin Tissue DNA kit and quantified using a Qubit fluorometer (Qubit dsDNA BR Assay Kit, Q32853, Invitrogen). Thirty-seven blood samples at T0 (pre-stress) and thirty-seven additional samples at T4 (post-stress) have been randomly selected for the downstream epigenomic analysis.

### Library preparation and sequencing

We followed the epiGBS protocol published by van Gurp et al [88]. As the method was developed for plants (with methylation occurring in CpG, CHH, and CHG context; H being any nucleotide but a cytosine) and used a single digestion approach, the protocol was modified to make it more suitable and straightforward for our vertebrate system. Particularly, a double digest instead of the single digest approach was implemented. We chose the restriction enzyme *MspI*, a standard choice in reduced representation bisulfite sequencing-like (RRBS) studies, as its recognition site targets CpG rich regions [204]. A second enzyme (*SfbI*) with a recognition site length of 8 bp was used to reduce fragment numbers, and thus to increase read depth per fragment. The choice of this enzyme was guided by *in silico* digestion of the European sea bass genome [76] using *simRAD* [205]. This genome is available at: https://www.ensembl.org/Dicentrarchus_labrax/Info/Index.

One single library was prepared for a set of 74 samples. For this library, 200 ng of DNA of each sample were digested in a 40 μl reaction, using 0.25 μl *MspI* (NEB, 20,000 U/ml R0106S), 0.25 μl *SbfI*-HiFi (NEB 20,000 U/ml R3642L) and 4 μl of 10X cutsmart buffer. The reaction was run overnight at 37°C. Unique forward and reverse adapter combinations allow multiplexing samples in the library. We added forward and reverse adapters in unique combinations (1 μl of adapter, 2.5 μM), 0.5 μl T4 Ligase (NEB, 400,000 U/ml m0202L), 6 μl of 10X T4 ligase buffer and 11.5μl water were added directly to the digested DNA. Sequences of adapters are provided as Additional File 1. Adapters were ligated for 3 h at 23°C followed by 10 min of enzyme inactivation at 65°C. After ligation, all samples were pooled and one third of the total volume was used in the following step. The mixture volume was reduced using a Qiaquick PCR purification kit (28104 Qiagen). The resulting product was cleaned a second time to ensure the removal of small fragments and adapter remnants using CleanPCR paramagnetic beads (Proteigene, CPCR-0050) with a ratio of 0.8X [sample: beads].

Since oligonucleotides were not phosphorylated (see Additional File 7), a nick translation was performed to repair the nick of the DNA at the restriction site and to fully methylate the hemi-methylated adapters. To do so, 19.25 μl of the concentrated and purified ligation pool were used in a 25 μl reaction, including 0.75 μl DNA Polymerase I (E. *coli*, 10,000 U/ml M0209S), 2.5 μl of 10 mM 5-Methylcytosine dNTP mix (Zymo Research, D1030), and incubated at 15 °C for one hour. The library was then treated with sodium bisulfite using the EZ DNA Methylation Gold Kit (D5005, Zymo Research) following manufacturer instructions to convert unmethylated cytosines to uracil, paying attention to the optimal DNA amount per reaction. After bisulfite conversion, 14 cycles of PCR (95°C for 1 min, 91°C for 10s, 65°C for 15s, 72°C for 10s and a final elongation step at 72°C for 5 min) were performed, followed by a paramagnetic bead clean up with a 0.8X ratio. The library quality (fragment size distribution, no adapters left, no primers left; fragment size range from 300 to 800 bp; see Additional File 8) was verified on an Agilent 5300 Fragment Analyzer (Santa Clara, USA). It was sequenced (paired-end, 150 bp) on one lane of a SP flow cell on an Illumina™ NovaSeq 6000 at the MGX sequencing facility in Montpellier, France.

### Bioinformatics analysis

Raw sequencing data were trimmed of low quality reads and adapter residues using *Trim Galore!* (v.0.6.4; available at https://www.bioinformatics.babraham.ac.uk/projects/trim_galore/). The trimmed reads were then demultiplexed using the `process_radtags` command of Stacks [206]. The option `disable_rad_check` was applied to avoid reads with bisulfite-modified restriction sites to be discarded. The demultiplexed reads were trimmed a second time on both 5’ and 3’ ends with the options `--clip_R1 --clip_R2` and `--three_prime_clip_R1 --three_prime_clip_R2` to remove introduced methylated cytosines during adapter ligation. Using the `bismark_genome_preparation` function of the program *bismark* [112], a bisulfite converted version of the European sea bass genome was prepared against which the demultiplexed and trimmed reads were mapped.

### Methylation analysis

The R package MethylKit [207] was used to determine differential methylation between pre- and post-stress fish. CpGs with less than 30 read depth and with coverage >99.9% of the distribution of read counts were filtered out to account for PCR bias. Coverage was normalized across samples, and only CpGs present in at least 20 out of 37 samples per group (56%) were kept for further analysis. Differentially methylated CpG sites were determined between pre- and post-stress individuals using logistic regression (`calculateDiffMeth` function). Pre-stress individuals were considered as the baseline. Cytosines were considered as differentially methylated when presenting at least 15% methylation difference (as in [83] for epigenomic variation in *D. labrax*), and a nominal *q*-value < 0.001 between pre- and post-stress fish. Reads presenting differentially methylated cytosines (DMCs) were extracted using Geneious (v.11.0; available at https://www.geneious.com/) and mapped along LGs of the European sea bass reference genome to produce a primary annotation of potential candidate genes. As the annotation of the European sea bass and the sea bream (*Sparus aurata*) reference genomes have been recently released on Ensembl (http://www.ensembl.org/index.html), gene names and models have been controlled, eventually including a TBLASTN search in case of discrepancy.

### Literature surveys and protein-protein association

A first literature survey was performed in order to decipher if genes found associated to European sea bass DMCs were candidates related to stress in vertebrates. We searched PubMed (https://pubmed.ncbi.nlm.nih.gov/) for studies in which they could have been mentioned as responding to stress. This approach was preferred to, e.g. gene ontology or enrichment analyses that are useful summaries of functional knowledge, but less suitable when searching how one or a specific set of genes might be involved more precisely in the stress response, if they are known to be associated to each other and/or how they were found related to stress (e.g. in which organism, context, study, organ or tissues). As relatively few DMCs were detected (see Results), this detailed approach was amenable.

The data mining considered a DMC relevant to the stress response when associated to a gene related to a stress phenotype (typically issued from genome-wide association studies, knock-out experiments or empirical evidence of impairment and dysfunction when facing stressors). Eight stress response themes were identified and used to structure the mining: ‘*blood*’ (including issues on red blood cells, blood homeostasis, blood volume, hemato- and erythropoiesis, platelets, hypo- and hypertension, angiogenesis), ‘*stress*’ (including stress-related genes, physiological disorders, HPI/HPA axis, adrenals, head kidney, stress resistance/response), ‘*immunity*’ (including issues on all types of leukocytes and immune response, cytokines, cytotoxicity and response to pathogens), ‘*hormones*’ (including issues related to catecholamines, corticosteroids, glucocorticoids, neuro- and peptide transmitters, other hormones related to the HPI axis in fish), ‘*brain*’ (including issues on hypothalamus and pituitary themselves, forebrain, limbic brain, neuronal development and plasticity, signaling pathways in the brain, serotonergic/dopaminergic activities), ‘*glucose*’ (including issues on glucose metabolism [systemic and brain], lactate, glycolysis, gluconeogenesis, [pro]insulin, diverse receptors, pancreatic β-cells), and ‘*behaviour*’ as stress-induced behavioural changes and not only physiological changes. Diverse health disorders related to stress often well-studied in mammalian models were screened, as well as some genetic disorders that affect behaviour. This may include changes in mood, emotional and affective states (e.g. fear, anxiety), cognitive traits (e.g. memory, learning abilities), or psychiatric disorders and neurodegenerative diseases (e.g. bipolar disorders, autism, schizophrenia). One additional category (‘*others*’) was included to report significant items known to be important for, e.g., epigenetic regulation that could not be directly addressed with previous themes. Cellular issues of stress such like, e.g., oxidative, chromatin or endoplasmic reticulum stress where not covered.

A second literature survey focused on the literature dealing with stress, stress response and high-throughput transcriptomics or epigenomics in fish. We searched PubMed for relevant articles over the period 2010-2020 (June) and, when available, their associated supplementary materials, and searching for difference in gene expression or methylation. Our goal was to evaluate if DMC-related genes detected here in RBCs are relevant to brain and/or processes specific to brain or if they are commonly detected whatever the target tissues and the nature of stressors. For this survey, we first mined gene expression studies that used blood within fish species in order to verify whether sea bass DMC-related genes were not commonly overexpressed in this tissue. In a second time, we focused on zebrafish for which functional annotation is well established. We searched for transcriptomics (RNA sequencing) studies dealing with stress in various tissues (liver, kidney, gill, brain, gonad, muscle, eye and skin). The following search string was used: ‘*zebrafish*’ AND ‘*RNA seq*\*’ AND ‘*stress*’ AND [‘*tissue name*’]. As mRNA studies often consider several tissues, results were manually curated. Studies whose title referred to single gene families (e.g. heat shock proteins) or microRNAs were discarded. Finally, fish epigenomic studies were also analyzed. About this issue, we first analyzed gene transcripts and differentially methylated genes in the single genome-wide epigenomic study performed so far in sea bass [83]. As genome-wide methylation studies remain relatively few in fish, this was extended to other species in a second time. Transcriptomics and methylation data sets that did not provide for gene names and/or gene symbols were discarded. Reported gene names and symbols were considered to be exact.

Finally, in order to complete these manually curated surveys, we used String (https://string-db.org/) to investigate if DMC-related genes encoded for proteins are known to interact together. We used our gene list as input and the zebrafish genome as a reference for annotation. This search was done without reference to the thematic stress categories mentioned above, and when one interaction was provided we specifically explored the literature for confirmation and relevant experimental evidence.

### Data analysis

A principal component analysis (PCA) on the methylation profiles of individual samples at DMCs was performed to analyse the potential grouping structures within our data set. Differences in the distribution of mean individual loading scores for pre- and post-stress fish were tested along each principal components axis using a Student *t*-test. Additionally, a hierarchical clustering was performed using Ward’s linkage method on Euclidean distances in order to explore whether sea bass families structure the data set.

## Supporting information

Captions and Additional Files except Add. File 3

## Abbreviations

ABLIM: actin-binding LIM kinase
AC: adenyl(-ate) cyclase
ACTH: adrenocorticotropic hormone
BDNF: Brain Derived Neurotrophic Factor
cAMP: cyclic adenosine monophosphate
CpG: cytosine-phosphate-guanine
CREB: cAMP response element-binding
CRH: corticotropin-releasing hormone
DMC: differentially methylated cytosine
DMR: differentially methylated region
epiGBS: epiGenotyping By sequencing
GDNF: glial cell line-derived neurotrophic factor
HPA: Hypothalamus-Pituitary-Adrenal
HPI: Hypothalamus-Pituitary-Interrenal
LG: linkage group
mRNA: messenger RNA
NCBI: National Center for Biotechnology Information
PACAP: Protein Adenylate Cyclase Activating Protein
POMC: pro-opiomelanocortin
RBC: red blood cell
RRBS: reduced representation bisulfite sequencing
UTR: untranslated region

## DECLARATIONS

### Ethics approvals and consent to participate

All experiments were performed in accordance with relevant guidelines and regulations. Nireus S.A. research facilities are certified and have obtained the codes for the rearing and use of fish for scientific purposes (EL04-BIOexp-01). All procedures on fish used in this study were approved by the Departmental Animal Care Committee following the Three Rs principle, in accordance with Greek (PD 56/2013) and EU (Directive 63/2010) legislation on the care and use of experimental animals. All sample procedures were performed by FELASA certified researchers (http://www.felasa.eu/).

### Consent for publication

Not applicable

### Availability of data and materials

Individual Illumina raw reads (one round of trimming) and processed files have been deposited in NCBI’s Gene Expression Omnibus [208] and are accessible through GEO Series accession number GSE153838 (https://www.ncbi.nlm.nih.gov/geo/query/acc.cgi?acc=GSE153838).

### Author contributions

MVK, ED, BG developed the protocol, MVK and ED conducted the lab work, EG ensured technical assistance and supervised sequencing. AS, AD, MP and CST established the study design used during the challenge test. Experiments with live fish were conducted by AD, AS, MP, with technical contribution of MVK. MVK performed data analyses under supervision of ED and BG. MVK, ED, BG drafted the manuscript; others authors reviewed drafts. CST and BG coordinated this study. All authors have read and approved the manuscript.

### Author details

See first page

## Acknowledgements

Data used in this work were partly produced through the GenSeq technical facilities at ISEM and benefited from the Montpellier Bioinformatics Biodiversity (MBB) facility, both supported by the LabEx CeMEB, and by ANR “Investissements d’Avenir” program (ANR-10-LABX-04-01). MGX acknowledges financial support from the France Génomique National infrastructure, also funded as part of “Investissement d’Avenir” program managed by ANR (ANR-10-INBS-09). We thank N. Wagemaker at the Radboud University (Nijmegen, The Netherlands) for his support, and Prof. C. Grunau at the University of Perpignan Via Domitia (France) for hosting M.V.K at various steps of this work. S. Ben Chehida helped during the last stages of manuscript preparation.

