## Supplementary material for "Genome-wide epigenomic stress response in the European sea bass (*Dicentrarchus labrax*, L.)": Captions and Additional Files except Add. File 3

**Genome scale epigenomic analysis of response to acute stress in the European sea bass (*Dicentrarchus labrax*, L.)**

**This file contains:**

**PAGE 3: Additional File 1:** Read numbers per sample and methylation patterns. Families, treatment (pre- *vs* post-stress) and sample codes are reported as in Fig. 3. The codes that will be used for individual deposits of samples within a NCBI sequence read archive (SRA) are given. Note that one single post-stress individual (T4_M17) retrieved a bisulfite conversion rate far lower than other individuals. It was kept for further analysis as mapping parameters were not found different from other individuals.

**PAGE 4: Additional File 2:** DMCs, DMRs and DMC-related genes. This material is one extended version of Table 1, providing details on, e.g. annotations and CpG context. Grey areas indicate DMCs grouped as DMRs.

**PAGE 5: Additional File 3:** Aim, rationales and details on the relationship of each sea bass DMC-related gene to stress according to the eight themes defied in the main text (see also Fig. 2). A short description is reported, some references are provided with doi or PMID identifiers. The table report high-throughput transcriptomic studies in which DMC-related genes detected in sea bass were found differentially expressed. The reference tissue is indicated and some key references provided at the bottom of the table.

**PAGE 7: Additional File 4:** List of zebrafish transcriptomic studies surveyed (*n* = 22). Results are reported per tissue. Genes found identical to our DMC-related genes found in sea bass are indicated for each study. Focal studies regarding brain and mentioned in the main text are indicated in red; list of genes pertaining to these studies are in Suppl. Mat. Table S4.

**PAGE 10: Additional file 5:** Summary of epigenomic studies (*n* = 20 out of 31 studied considered) that reported the DMC-related genes found in the European sea bass in stress - or other – experiments as differentially methylated. Species, tissues and genes detected in each study are indicated. Tissues that reported no match are not reported. The six studies that considered the brain tissue presented at least one gene found differentially methylated in red blood cells of sea bass. See main text for further comments. Studies surveyed are provided after the table.

**PAGE 13: Additional File 6:** Output of the String search using the DMC-related genes as input. Eight associations are shown and few discussed in the text as relevant. Note that the search was made using annotation on zebrafish *Danio rerio* and gene symbols might be different than those reported in tables. String automatically changes annotations to report only those used in the reference zebrafish genome. The correspondence is reported for the most significant (*CRTC2, NOL4LB, SASH1A*). *GLG1* and *CSMD3a* have no annotation in *D. rerio* genome and are not reported. Colored traits at each association have different meanings that are not relevant to (e.g. literature search, experimental evidence), but that have all been explored in this study.

**PAGE 14: Additional File 7:** Adapter sequences used in this study, X = methylated cytosine, C = non methylated cytosines. Non methylated cytosines are replaced during nick translation with m-dCTPs and therefore a fully methylated adapter is reconstituted.

**PAGE 15: Additional File 8:** Fragment size distribution of the epiGBS library sequenced in this study.

**Additional File 1**

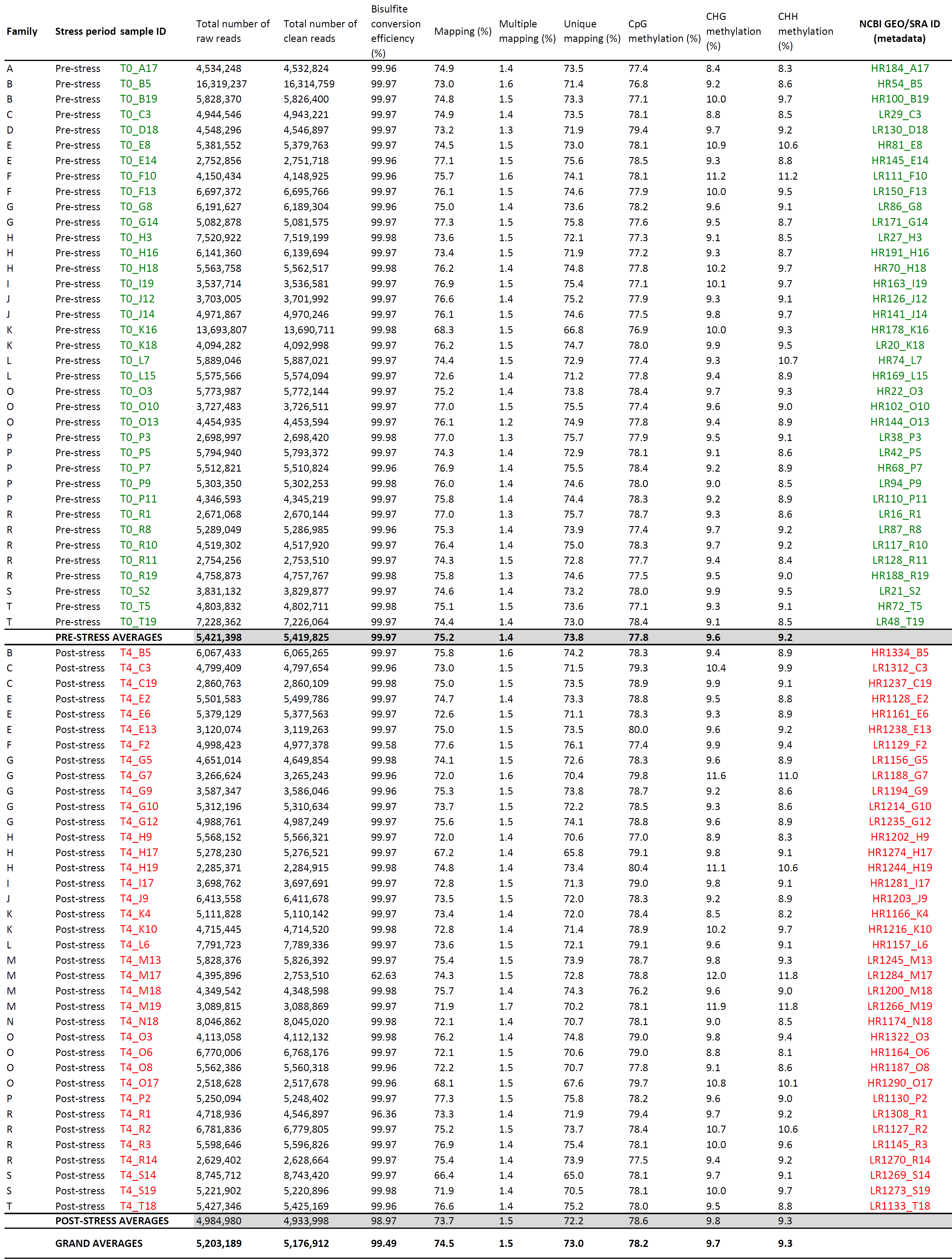

**Additional File 2**

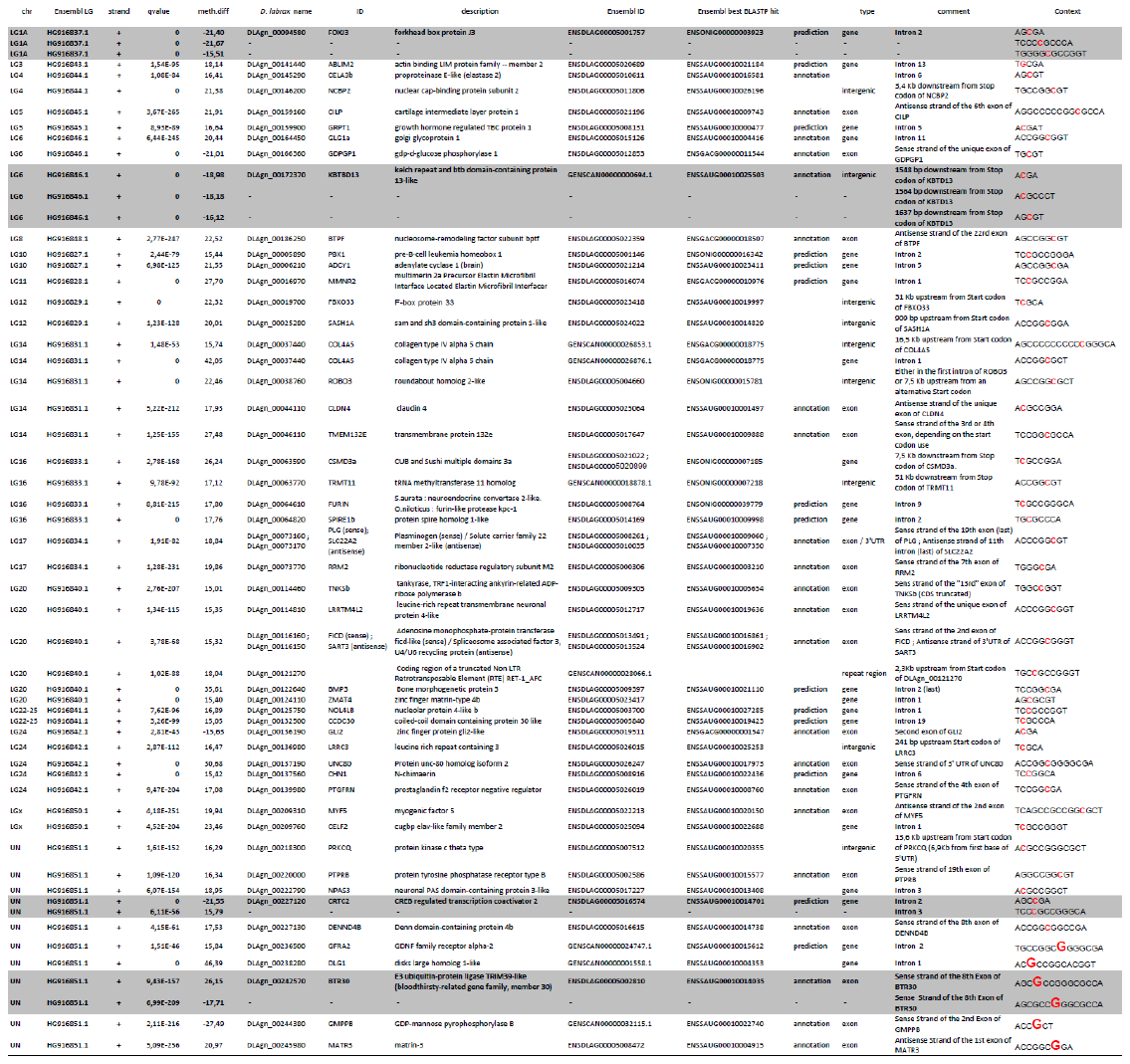

**Additional File 3**

**Available as a separate additional file, downloaded separately or available upon request**

**Caption:**

**Aim:** This Table provides summaries of our literature survey on 51 DMC-related genes reported after genome-wide epiGBS study of red blood cells RBCs) of 70 pre- and post-stress sea bass (74 samples). For each DMC-related gene, search was associated to eight search categories that represent one integrative view of the stress response in fish: ‘blood’, 'stress’, ‘immunity’, ‘hormones’, ‘brain’, ‘glucose’, ‘behavior’, and ‘others’ when necessary. Description of these themes is reported in the main text of the paper and a graphical expression is given in Fig. 2 of the manuscript. It reports these categories for each DMC-related gene as in Figure 2, provides a short written summary for each gene and associated references with their doi or PMID identifier.

**Rationale:**

**1 -** Genes implicated in a DMR are highlighted in grey; genes with a DMC within the gene body are in red, while genes found close to one intergenic DMC are given in black (details on exact location of DMCs like exon, intron and distance to the gene are reported in Table 1 of the main text as well as in Supplementary Materials Table S1).

**2** - Situations discussed in the main text are NOT reproduced there (i.e. specific DMC-related genes connected to BDNF (Brain-Derived Neurotrophic Factor). *IF* these genes are implied in other aspects of the stress response, additional information is provided in the table.

**3 - SEA BASS:** Genes also found differentially methylated or differentially transcribed in Anastasiadi and Piferrrer (2019) are indicated **(AP)** and the tissue in which differential expression/methylation was recorded (liver, muscle, testis for transcriptomics, these three tissues + brain for the CpG analyses).

**4** – As transcriptomic publications report DMC-related genes (see main text), genes concerned by these studies are indicated in each case by the initials of the first author (see below). Red indicate zebrafish studies, while black and blue are for a sea bass and a *Betta splendens* (Siamese fighting fish) study, respectively.

References are:

**(R):** Rey S., Boltana S., Vargas R., Roher N., Mackenzie S. 2013. Combining animal personalities with transcriptomics resolves individual variation within a wild-type zebrafish population and identifies underpinning molecular differences in brain function. Mol. Ecol. 22: 6100–6115. doi: 10.1111/mec.12556

**(W):** Wong R.Y., Lamm M.S., Godwin J. 2015. Characterizing the neurotranscriptomic states in alternative stress coping styles. BMC Genomics 16(1): 425. doi: 10.1186/s12864-015-1626-x

**(GP):** Greenwood A.K., Peichel C.L. 2015. Social regulation of gene expression in threespine sticklebacks. PLoS One 10(9): e0137726. doi:10.1371/journal.pone.0137726

**(B):** Boswell M., Lu Y., Boswell W., Savage M., Hildreth K., Salinas R., Walter C.A., Walter R.B. 2019. Fluorescent Light Incites a Conserved Immune and Inflammatory Genetic Response within Vertebrate Organs (*Danio rerio, Oryzias latipes* and *Mus musculus*). Genes 10(4): 271. doi: 10.3390/genes10040271

**(O):** Oliveira RF, Simões JM, Teles MC, Oliveira CR, Becker JD, Lopes JS. 2016. Assessment of fight outcome is needed to activate socially driven transcriptional changes in the zebrafish brain. Proc. Natl Acad. Sci. USA 113(5):E654-E661. doi:10.1073/pnas.1514292113

**(T):** Thyme S.B., Pieper L.M., Li E.H., Pandey S., Wang Y., Morris N.S., Sha C., Choi J.W., Herrera K.J., Soucy E.R., Zimmerman S., Randlett O., Greenwood J., McCarroll S.A., Schier A.F. 2019. Phenotypic landscape of schizophrenia-associated genes defines candidates and their shared functions. Cell 177(2): 478-491.e20. doi: 10.1016/j.cell.2019.01.048.

**(Wa):** Wang M., Xu G., Tang Y. et al. 2020. Transcriptome analysis of the brain provides insights into the regulatory mechanism for *Coilia nasus* migration. BMC Genomics 21: 410. doi: 10.1186/s12864-020-06816-3

**(CP):** Chaves-Pozo E, Bandín I, Olveira JG, et al. 2019. European sea bass brain DLB-1 cell line is susceptible to nodavirus: A transcriptomic study. Fish Shellfish Immunol. 86: 14‐24. doi:10.1016/j.fsi.2018.11.024

**(V):** Vu TD, Iwasaki Y, Shigenobu S, et al. 2020. Behavioral and brain- transcriptomic synchronization between the two opponents of a fighting pair of the fish *Betta splendens*. *PLoS Genet*. 16(6):e1008831. doi:10.1371/journal.pgen.1008831

**NOTE** that when DMC-related genes are associated to **(W)**, **(GP)** or **(R)** studies that dominate reports of DMC-related genes (Suppl. Mat. Text S1), no description is provided for 'brain', 'stress', 'behavior'. ONLY descriptions and additional references (+ doi) for the other themes –sometimes fish species - are reported.

**5 -** Cells highlighted in blue indicate DMR-related genes reported ONLY in studies cited in the second column of the table. No additional or relevant references were found.

**6** - Studies relevant to a 'stress context' or 'stress environment' - including developmental diseases, knock-out experiments - are referenced with doi for each gene. Some general references might be reported.

**Additional File 4**

**Transcriptomics Survey**

**BRAIN (total number of distinct DMC-related genes found in common with our sea bass study in brain: *38,* details in Fig. 2 and Suppl. Mat. Table S4)**

**1**: => **CSMD3** => Thyme SB, Pieper LM, Li EH, Pandey S, Wang Y, Morris NS, Sha C, Choi JW, Herrera KJ, Soucy ER, Zimmerman S, Randlett O, Greenwood J, McCarroll SA, Schier AF. Phenotypic Landscape of Schizophrenia-Associated Genes Defines Candidates and Their Shared Functions. Cell. 2019 177(2):478-491.e20. doi: 10.1016/j.cell.2019.01.048

**2**: => **DENND4B, CLDN4** => Boswell M, Lu Y, Boswell W, Savage M, Hildreth K, Salinas R, Walter CA, Walter RB. Fluorescent Light Incites a Conserved Immune and Inflammatory Genetic Response within Vertebrate Organs (*Danio rerio, Oryzias Latipes* and *Mus Musculus*). Genes (Basel). 2019 10(4):271. doi: 10.3390/genes10040271

**3:** **=>** **FOXJ3, ABLIM2, NCBP2, GLG1A, ADCY1b, PBX1, SASH1A, COL4A5, ROBO3, CSMD3, FURIN, SPIRE1b, TRMT11, SLC22A2, RRM2,** **BMP3, FICD, SART3, TNKS**, **ZMAT4b, CCDC30, CHN1, GLI2**, **PTGFRN, UNC80, CELF2, DENND4B, CRTC2, GFRA2,** **GMPPB, MATR3, NPAS3, PRKCQ, PTPRB** (**N = 34** ; see Supp. Mat Table S4) => Wong RY, Lamm MS, Godwin J. Characterizing the neurotranscriptomic states in alternative stress coping styles. BMC Genomics. 2015 16(1):425. doi: 10.1186/s12864-015-1626-x

**4:** **=> none** => Porseryd T, Volkova K, Reyhanian Caspillo N, Källman T, Dinnetz P, Porsh Hällström I. Persistent Effects of Developmental Exposure to 17α-Ethinylestradiol on the Zebrafish (*Danio rerio*) Brain Transcriptome and Behavior. Front Behav Neurosci. 2017 11:69. doi: 10.3389/fnbeh.2017.00069

**5:** **=> none** => Wong RY, Oxendine SE, Godwin J. Behavioral and neurogenomic transcriptome changes in wild-derived zebrafish with fluoxetine treatment. BMC Genomics. 2013 14:348. doi: 10.1186/1471-2164-14-348

**6:** **=> no data at the gene level** => Li Z, Li W, Zha J, Chen H, Martyniuk CJ, Liang X. Transcriptome analysis reveals benzotriazole ultraviolet stabilizers regulate networks related to inflammation in juvenile zebrafish (Danio rerio) brain. Environ Toxicol. 2019 34(2):112-122. doi: 10.1002/tox.22663

**7**: **=> DLG1, UNC80, PTGFRN, NOL4B, TNKS,** **ROBO3, COL4A5, SASH1A, ADCY1b**, **GLG1a** (**N= 10** ; see Supp. Mat Table S4) => Greenwood AK, Peichel CL. Social Regulation of Gene Expression in Threespine Sticklebacks. PLoS One. 2015 10(9):e0137726. doi: 10.1371/journal.pone.0137726

**8**: **=>** **NCBP2, CILP1, FURIN, RRM2, GLI2, DLG1, CELF2** (**N= 7** ; see Supp. Mat Table S4) => Rey S., Boltana S., Vargas R., Roher N., Mackenzie S. 2013. Combining animal personalities with transcriptomics resolves individual variation within a wild-type zebrafish population and identifies underpinning molecular differences in brain function. Mol. Ecol. 22: 6100–6115 doi: 10.1111/mec.12556

**9 => CLDN** => Wang M., Xu G., Tang Y. et al. 2020. Transcriptome analysis of the brain provides insights into the regulatory mechanism for *Coilia nasus* migration. BMC Genomics 21: 410. doi: 10.1186/s12864-020-06816-3

**10 => DLG1** => Oliveira RF, Simões JM, Teles MC, Oliveira CR, Becker JD, Lopes JS. 2016. Assessment of fight outcome is needed to activate socially driven transcriptional changes in the zebrafish brain. Proc. Natl Acad. Sci. USA 113(5):E654-E661. doi:10.1073/pnas.1514292113

**TOTAL NUMBER OF DMC-RELATED GENES FOUNDS IN COMMON WITH OUR EUROPEAN SEA BASS STUDY FOR TISSUES (BRAIN EXCLUDED) IS: 15 (20 occurrences, 15 distinct)**

**LIVER** (total number of DMC-related genes found in common with our sea bass study in the liver: **6**)

**1: => none** (only GO-terms and KEGG pathways, few candidate genes used, other not reported) => Zheng M, Lu J, Zhao D. Toxicity and Transcriptome Sequencing (RNA-seq) Analyses of Adult Zebrafish in Response to Exposure Carboxymethyl Cellulose Stabilized Iron Sulfide Nanoparticles. Sci Rep. 2018 ;8(1):8083. doi: 10.1038/s41598-018-26499-x

**2: => ROBO3, NCBP2** => Dhanasiri AK, Fernandes JM, Kiron V. Liver transcriptome changes in zebrafish during acclimation to transport-associated stress. PLoS One. 2013 8(6):e65028. doi: 10.1371/journal.pone.0065028

**3: => PTPRB** => Boswell M, Lu Y, Boswell W, Savage M, Hildreth K, Salinas R, Walter CA, Walter RB. Fluorescent Light Incites a Conserved Immune and Inflammatory Genetic Response within Vertebrate Organs (*Danio rerio, Oryzias Latipes* and *Mus Musculus*).Genes (Basel). 2019 10(4):271. doi: 10.3390/genes10040271

**4: => COL4A5** (but full list of differentially expressed genes not provided) => Zhang J, Liu L, Ren L, Feng W, Lv P, Wu W, Yan Y. The single and joint toxicity effects of chlorpyrifos and beta-cypermethrin in zebrafish (*Danio rerio*) early life stages. J Hazard Mater. 2017 334:121-131. doi: 10.1016/j.jhazmat.2017.03.055

**5: => GLI2** => Xu H, Lam SH, Shen Y, Gong Z. Genome-wide identification of molecular pathways and biomarkers in response to arsenic exposure in zebrafish liver. PLoS One. 2013 8(7): e68737. doi: 10.1371/journal.pone.0068737

**6: => BTR30** => Kang Q, Hu M, Jia J, Bai X, Liu C, Wu Z, Chen W, Li M. Global Transcriptomic Analysis of Zebrafish Glucagon Receptor Mutant Reveals Its Regulated Metabolic Network. Int J Mol Sci. 2020 21(3):724. doi: 10.3390/ijms21030724

**7: => none** (but full list of differentially expressed genes not provided) => Vergauwen L, Benoot D, Blust R, Knapen D. Long-term warm or cold acclimation elicits a specific transcriptional response and affects energy metabolism in zebrafish. Comp Biochem Physiol A Mol Integr Physiol. 2010 157(2):149-57. doi: 10.1016/j.cbpa.2010.06.160

______________

**MUSCLE** (total number of DMC-related genes found in common with our sea bass study in the muscle: **0**)

**1: => none** (also for EYE) Arcanjo C, Armant O, Floriani M, Cavalie I, Camilleri V, Simon O, Orjollet D, Adam-Guillermin C, Gagnaire B. Tritiated water exposure disrupts myofibril structure and induces mis-regulation of eye opacity and DNA repair genes in zebrafish early life stages. Aquat Toxicol. 2018 200:114-126. doi: 10.1016/j.aquatox.2018.04.012

**2: => none** => Cambier S, Gonzalez P, Durrieu G, Maury-Brachet R, Boudou A, Bourdineaud JP. Serial analysis of gene expression in the skeletal muscles of zebrafish fed with a methylmercury-contaminated diet. Environ Sci Technol. 2010 44(1):469-75. doi: 10.1021/es901980t

______________

**GONADS** (total number of DMC-related genes found in common with our sea bass study in the gonads: **3**)

**1: => ROBO3, RRM2, CELA3** => Porseryd T, Reyhanian Caspillo N, Volkova K, Elabbas L, Källman T, Dinnétz P, Olsson PE, Porsch-Hällström I. Testis transcriptome alterations in zebrafish (*Danio rerio*) with reduced fertility due to developmental exposure to 17α-ethinyl estradiol. Gen Comp Endocrinol. 2018 262:44-58. doi:10.1016/j.ygcen.2018.03.011

______________

**INTESTINE** (total number of DMC-related genes found in common with our sea bass study in the intestine: **6**)

**1**: **=> CELA3, SLC22A2, BPTF, DENND4B, CELF2, GLG1a** => Zhang QL, Dong ZX, Luo ZW, Zhang M, Deng XY, Guo J, Wang F, Lin LB. The impact of mercury on the genome-wide transcription profile of zebrafish intestine. J Hazard Mater. 2020 389:121842. doi: 10.1016/j.jhazmat.2019.121842

______________

**SKIN** (total number of DMC-related genes found in common with our sea bass study in the skin: **5**)

**1: => RRM2, COL4A5, GFRA2, KBTBD13, GLI2** => Boswell M, Lu Y, Boswell W, Savage M, Hildreth K, Salinas R, Walter CA, Walter RB. Fluorescent Light Incites a Conserved Immune and Inflammatory Genetic Response within Vertebrate Organs (*Danio rerio, Oryzias Latipes* and *Mus Musculus*).Genes (Basel). 2019 10(4): 271. doi: 10.3390/genes10040271

**Additional File 5**

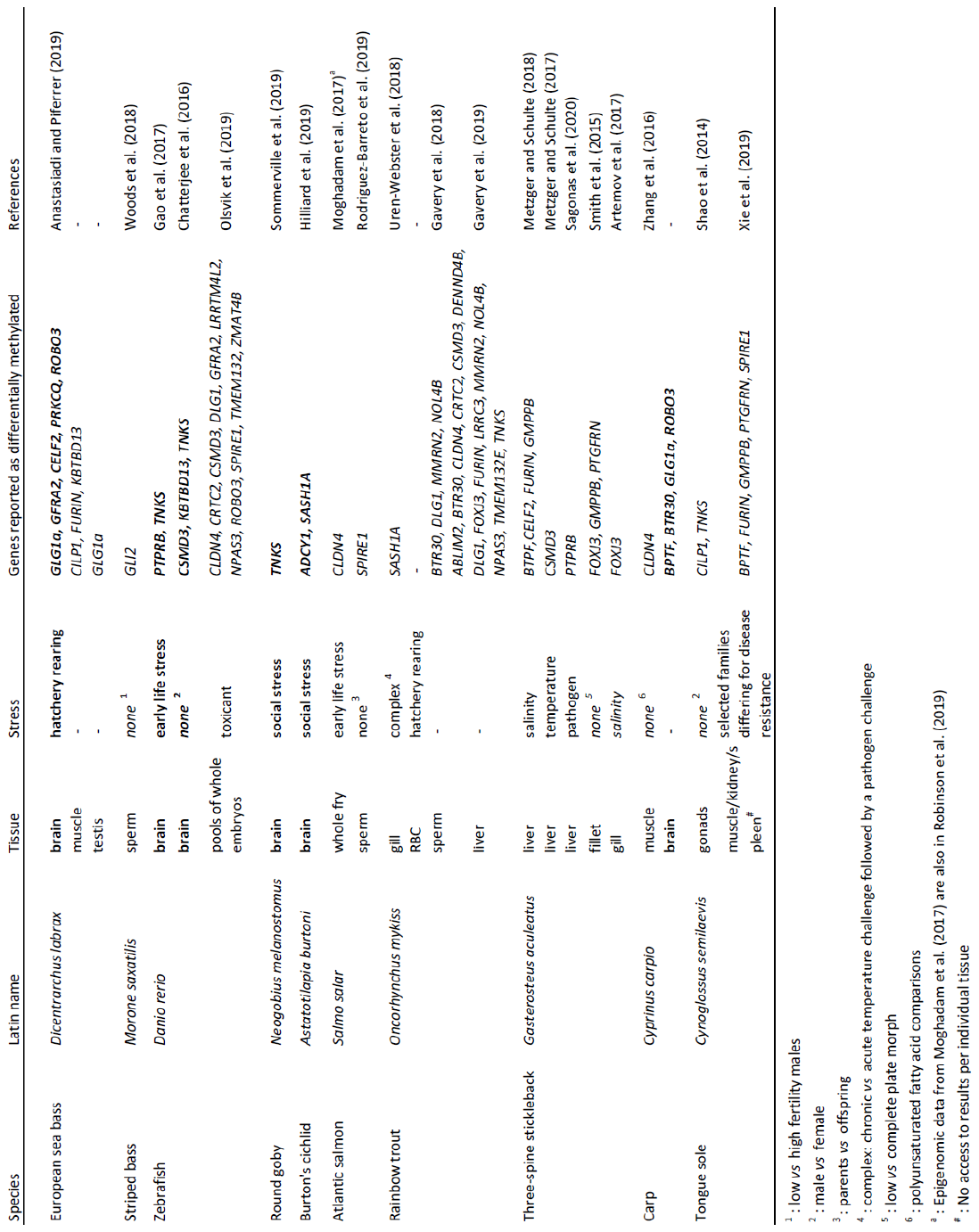

References of epigenome-wide studies considered in the epigenomic survey (alphabetical order). Studies not mentioned in the table above retrieved no match with the DMC-related genes detected in our European sea bass study.

1 - Anastasiadi D., Piferrer F. 2019. Epimutations in developmental genes underlie the onset of domestication in farmed European sea bass. *Mol. Biol. Evol*. 36(10):2252-2264. doi: 10.1093/molbev/msz153

2 - Artemov A.V., Mugue N.S., Rastorguev S.M., Zhenilo S., Mazur A.M., Tsygankova S.V., Boulygina E.S., Kaplun D., Nedoluzhko A.V., Medvedeva Y.A., et al. 2017. Genome-wide DNA methylation profiling reveals epigenetic adaptation of stickleback to marine and freshwater conditions. *Mol. Biol. Evol*. 34(9): 2203–2213. doi: 10.1093/molbev/msx156

3 - Baerwald M.R., Meek M.H., Stephens M.R. et al. 2016. Migration-related phenotypic divergence is associated with epigenetic modifications in rainbow trout. *Mol Ecol*. 25(8): 1785‐1800. doi: 10.1111/mec.13231

4 - Berbel-Filho WM, Berry N, Rodríguez-Barreto D, Rodrigues Teixeira S, Garcia de Leaniz C, Consuegra S. 2020. Environmental enrichment induces intergenerational behavioural and epigenetic effects on fish*. Mol Ecol.* doi:10.1111/mec.15481

5 - Boltana S., Aguilar A., Sanhueza N. et al. 2018. Behavioral fever drives epigenetic modulation of the immune response in fish. *Front. Immunol*. 9: 1241. doi: 10.3389/fimmu.2018.01241

6 - Chatterjee A., Lagisz M., Rodger E.J. et al. 2016. Sex differences in DNA methylation and expression in zebrafish brain: a test of an extended 'male sex drive' hypothesis. *Gene* 590(2): 307‐316. doi:10.1016/j.gene.2016.05.042

7 - Gao D., Wang C., Xi Z., Zhou Y., Wang Y., Zuo Z. 2017. Early-life benzo[a]pyrene exposure causes neurodegenerative syndromes in adult zebrafish (*Danio rerio*) and the mechanism involved. *Toxicol. Sci.* 157(1): 74‐84. doi: 10.1093/toxsci/kfx028

8 - Gavery M.R., Nichols K.M., Goetz G.G., Middleton M.A., Swanson P. 2018. Characterization of genetic and epigenetic variation in sperm and red blood cells from adult hatchery and natural-origin steelhead, *Oncorhynchus mykiss. G3: Genes, Genomes, Genetics* 8(11): 3723-3736. doi: 10.1534/g3.118.200458

9 - Gavery, M.R.; Nichols, K.M.; Berejikian, B.A.; Tatara, C.P.; Goetz, G.W.; Dickey, J.T.; Van Doornik, D.M.; Swanson, P. 2019. Temporal dynamics of DNA methylation patterns in response to rearing juvenile steelhead (*Oncorhynchus mykiss*) in a hatchery versus simulated stream environment. Genes 10: 356. doi: 10.3390/genes10050356

10 - Heckwolf M.J., Meyer B.S., Häsler R., Höppner M.P., Eizaguirre C., Reusch T.B.H. 2020. Two different epigenetic information channels in wild three-spined sticklebacks are involved in salinity adaptation. *Sci. Adv*. 6(12): eaaz1138. doi:10.1126/sciadv.aaz1138

11 - Hilliard A.T., Xie D., Ma Z., Snyder M.P., Fernald R.D. 2019. Genome-wide effects of social status on DNA methylation in the brain of a cichlid fish, *Astatotilapia burtoni*. *BMC Genomics* 20(1): 699. doi: 10.1186/s12864-019-6047-9

12 - Hu J., Pérez-Jvostov F., Blondel L., Barrett R.D.H. 2018. Genome-wide DNA methylation signatures of infection status in Trinidadian guppies (*Poecilia reticulata*). *Mol. Ecol*. 27(15): 3087–3102. doi: 10.1111/mec.14771

13 - Laporte M., Le Luyer J., Rougeux C., Dion-Côté A.M., Krick M., Bernatchez L. 2019. DNA methylation reprogramming, TE derepression, and postzygotic isolation of nascent animal species. *Sci. Adv.* 5(10): eaaw1644. doi: 10.1126/sciadv.aaw1644

14 - Le Luyer J., Laporte M., Beacham T.D., Kaukinen K.H., Withler R.E., Leong J.S., Rondeau E.R., Koop B.F., Bernatchez L. 2017. Hatchery-induced epigenetic modification in salmon. *Proc. Natl. Acad. Sci. USA* 114(49): 12964-12969. doi: 10.1073/pnas.1711229114

15 - Metzger D.C.H., Schulte P.M. 2017. Persistent and plastic effects of temperature on DNA methylation across the genome of threespine stickleback (*Gasterosteus aculeatus*). *Proc. Biol. Sci*. 284(1864): 20171667. doi:10.1098/rspb.2017.1667

16 - Metzger D.C.H., Schulte P.M. 2018. The DNA methylation landscape of stickleback reveals patterns of sex chromosome evolution and effects of environmental salinity. *Genome Biol. Evol*. 10(3): 775‐785. doi:10.1093/gbe/evy034

17 - Moghadam H.K., Johnsen H., Robinson N. et al. 2017. Impacts of early life stress on the methylome and transcriptome of Atlantic salmon. *Sci. Rep*. 7: 5023 doi: 10.1038/s41598-017-05222-2

18 - Olsvik PA, Whatmore P, Penglase SJ, Skjærven KH, Anglès d'Auriac M, Ellingsen S. 2019. Associations between behavioral effects of bisphenol A and DNA methylation in zebrafish embryos. *Front. Genet*. 10: 184. doi: 10.3389/fgene.2019.00184

19 - Robinson N.A., Johnsen H., Moghadam H., Andersen Ø., Tveiten H. 2019. Early developmental stress affects subsequent gene expression response to an acute stress in Atlantic salmon: an approach for creating robust fish for aquaculture? *G3: Genes, Genomes, Genetics* 9(5): 1597-1611. doi: 10.1534/g3.119.400152

20 - Rodriguez Barreto D., Garcia de Leaniz C., Verspoor E., Sobolewska H., Coulson M., Consuegra S. 2019. DNA methylation changes in the sperm of captive-reared fish: A route to epigenetic introgression in wild populations. *Mol Biol Evol*. 36(10): 2205‐2211. doi: 10.1093/molbev/msz135

21 - Sagonas K., Meyer B.S., Kaufmann J., Lenz T.L., Häsler R., Eizaguirre C. 2020. Experimental parasite infection causes genome-wide changes in DNA methylation. *Mol. Biol. Evol*. doi: 10.1093/molbev/msaa084

22 - Shao C., Li Q., Chen S. et al. 2014. Epigenetic modification and inheritance in sexual reversal of fish. *Genome Res*. 24(4):604-615. doi: 10.1101/gr.162172.113

23 - Smith G., Smith C., Kenny J.G., Chaudhuri R.R., Ritchie M.G. 2015. Genomewide DNA methylation patterns in wild samples of two morphotypes of threespine stickleback (*Gasterosteus aculeatus*). *Mol. Biol. Evol*. 32(4): 888–895. doi: 10.1093/molbev/msu344

24 - Somerville V., Schwaiger M., Hirsch P.E. et al. 2019. DNA methylation patterns in the round goby hypothalamus support an on-the-spot decision scenario for territorial behavior. *Genes* 10(3): 219. doi: 10.3390/genes10030219

25 - Todd E.V., Ortega-Recalde O., Liu H. et al. 2019. Stress, novel sex genes, and epigenetic reprogramming orchestrate socially controlled sex change. *Sci. Adv*. 5(7): eaaw7006. doi: 10.1126/sciadv.aaw7006

26 - Uren Webster T.M., Rodriguez-Barreto D., Martin S.A.M., Van Oosterhout C., Orozco-terWengel P., Cable J. et al. 2018. Contrasting effects of acute and chronic stress on the transcriptome, epigenome, and immune response of Atlantic salmon. *Epigenetics* 13(12):, 1191-1207. doi: 10.1080/15592294.2018.1554520

27 - Wan Z.Y., Xia J.H., Lin G., Wang L., Lin V.C.L., Yue G.H. 2016. Genome-wide methylation analysis identified sexually dimorphic methylated regions in hybrid tilapia. *Sci. Rep.* 6: 35903. doi: 10.1038/srep35903

28 - Woods III, L.C., Li Y., Ding Y. et al. 2018. DNA methylation profiles correlated to striped bass sperm fertility. *BMC Genomics* 19: 244. doi: 10.1186/s12864-018-4548-6

29 - Xiu Y., Shao C., Zhu Y., Li Y., Gan T., Xu W., Piferrer F., Chen S. 2019. Differences in DNA methylation between disease-resistant and disease-susceptible Chinese tongue sole (*Cynoglossus semilaevis*) families. *Front. Genet.* 10: 847. doi: 10.3389/fgene.2019.00847

30 - Zhang C., Hoshida Y., Sadler K.C. 2016. Comparative epigenomic profiling of the DNA methylome in mouse and zebrafish uncovers high interspecies divergence. *Front. Genet*. 7: 110. doi: 10.3389/fgene.2016.00110

31 - Zhang H., Xu P., Jiang Y., Zhao Z., Feng J., Tai R., Dong C., Xu J. 2019. Genomic, transcriptomic, and epigenomic features differentiate genes that are relevant for muscular polyunsaturated fatty acids in the common carp. *Front. Genet*. 10: 217. doi: 10.3389/fgene.2019.00217

**Additional File 6**

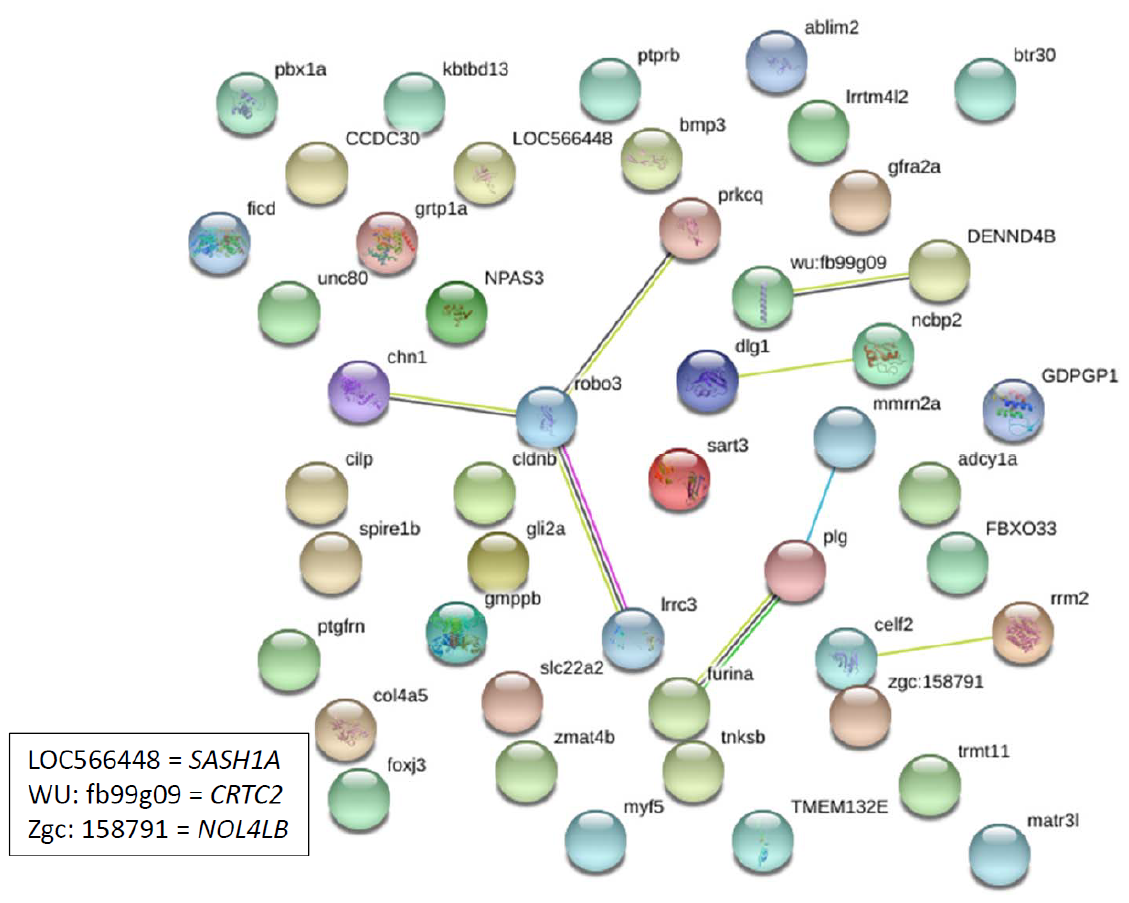

**Additional File 7**

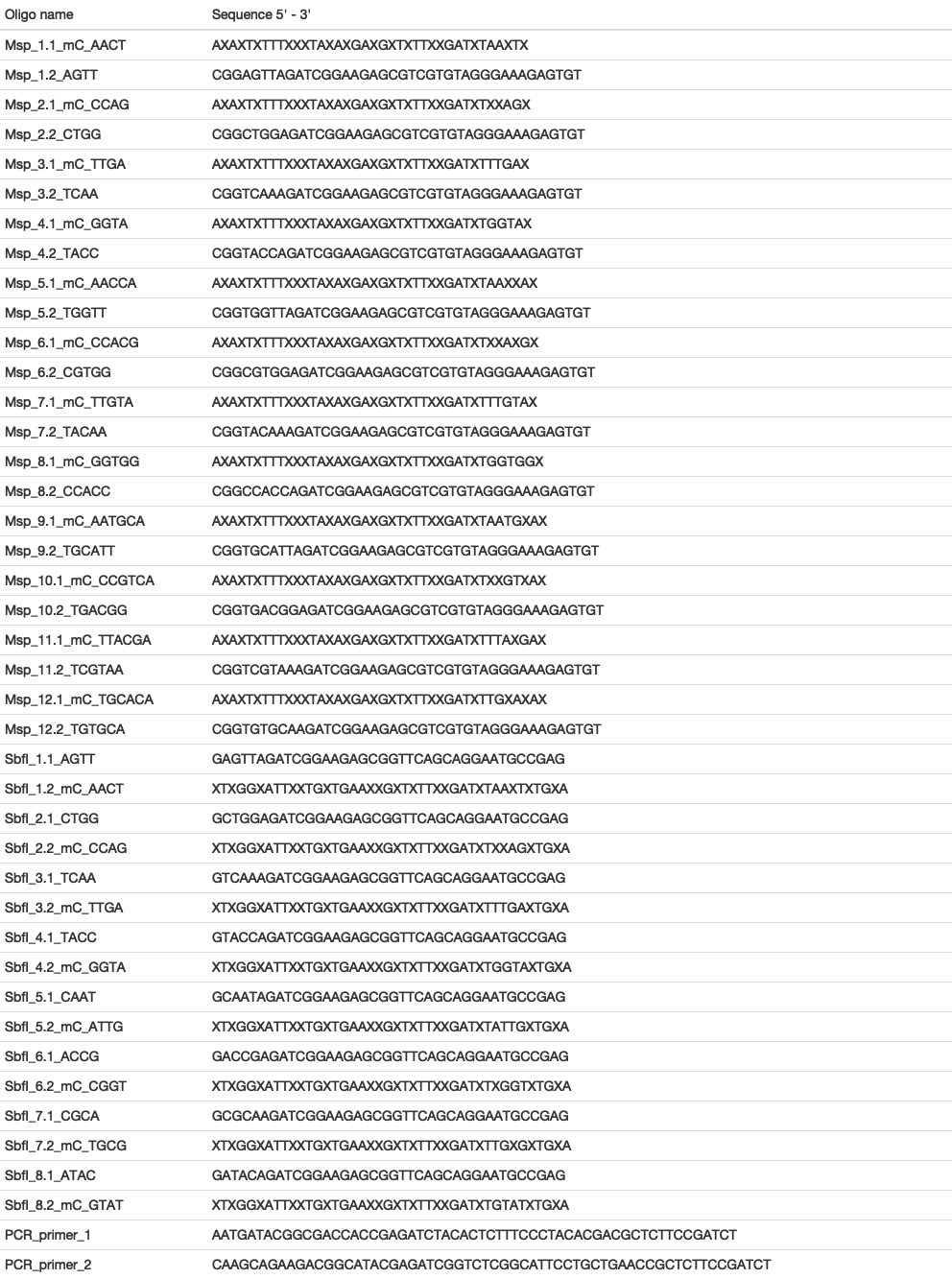

**Additional File 8**

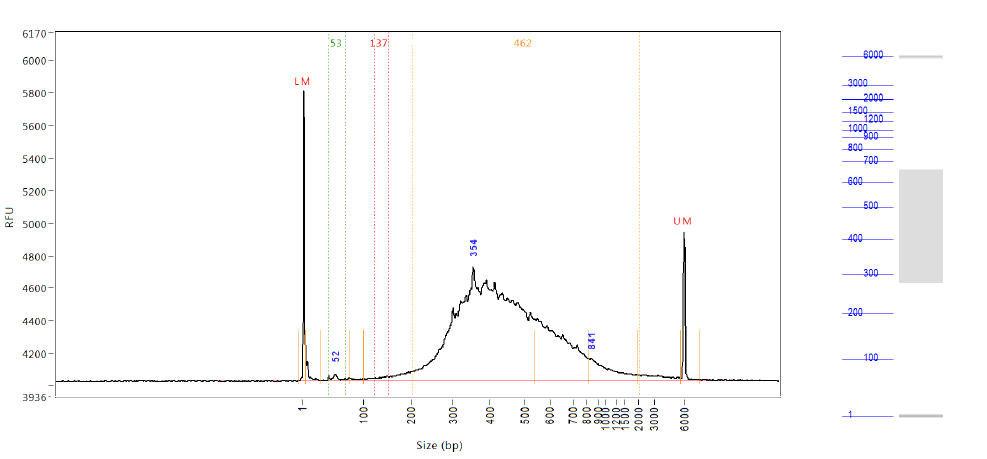
